# The Flowering Hormone Florigen Accelerates Secondary Cell Wall Biogenesis to Harmonize Vascular Maturation with Reproductive Development

**DOI:** 10.1101/476028

**Authors:** Akiva Shalit-Kaneh, Tamar Eviatar–Ribak, Guy Horev, Naomi Suss, Roni Aloni, Yuval Eshed, Eliezer Lifschitz

**Affiliations:** Department of Biology, Technion IIT; Lorey I. Lokey Center for Life Sciences & Engineering, Technion; School of Plant Sciences and Food Security, Tel Aviv University, Tel Aviv 69978, Israel; Department of Plant and Environmental Sciences Weizmann Institute of Science, Rehovot, Israel

## Abstract

The protein hormone florigen is a universal systemic inducer of flowering and a generic growth terminator across meristems. To understand the developmental rational for its pleiotropic functions and to uncover the deep cellular systems mobilized by florigen beyond flowering we explored termination of radial expansion of stems. Employing the power of tomato genetics along with RNAseq and histological validations we show that endogenous, mobile, or induced florigen accelerates secondary cell wall biogenesis (SCWB), and hence vascular maturation, independently of flowering. This finding is supported by a systemic florigen antagonist from the non-flowering *Ginkgo biloba*, which arrests SCWB and by *MADS* and *MIF* genes downstream of florigen that similarly suppress or enhance, respectively, vascular maturation independent of flowering. We also show that florigen is remarkably stable and distributed to all organs regardless of existing endogenous levels. By accelerating SCWB, florigen reprograms the distribution of resources, signals and mechanical loads required for the ensuing reproductive phase it had originally set into motion.

**Developmental Highlights:**

- Florigen accelerates SCWB: A prime case for a long-range regulation of a complete metabolic network by a plant hormone.
- The dual acceleration of flowering and vascular maturation by Florigen provides a paradigm for a dynamic regulation of global, independent, developmental programs.
- The growth termination functions of florigen and the auto-regulatory mechanism for its production and distribution provide a communication network enveloping the shoot system.
- A stable florigen provides a possible mechanism for the quantitative regulation of flowering
- Lateral stimulation of xylem differentiation links the phloem-travelling florigen with the annual rings in trunks.
- MADS genes are common relay partners in Florigen circuits; vascular maturation in stems and reproductive transition in apical meristems.

## Introduction

Nine decades ago, innovative grafting experiments firmly established a hypothetical signal, dubbed florigen, as a universal systemic inducer of flowering, produced in leaves and transported to the apical meristems (Chailakhyan, 1936, Zeevaart, 1976). However, the discovery of flowering pathways in Arabidopsis, which predominantly converge on the gene *FT* (Koornneef et al., 1991, Simpson and Dean, 2002), practically sent the elusive florigen into oblivion. A surprising twist in the odyssey of florigen emerged with the establishment in tomato of a one-to-one genetic relationship between florigen and the *SINGLE FLOWER TRUSS, SFT* gene, an ortholog of *FT* (Lifschitz et al., 2006, Sparks et al., 2013, Yung et al, 2007, Kobayashi and Weigel, 2007). This solitary genetic origin, unique among plant hormones, instituted florigen as a protein hormone universally encoded by *FT* orthologs, thus resolving the apparently conflicting paradigms.

More recent species-wide observations showed that in addition to flowering, florigen induces a plethora of morphogenetic effects all of which may be traced back to growth terminations: inactivation of *SFT* in tomato stimulates lateral leaf meristems but its overexpression converted compound leaves into simple leaves and restricted radial expansion, generating slender stems (Fig. 1B and 1C). Florigenic signals in addition regulate tuberization in potato, leaf size in Maryland Mammoth tobacco, cluster shape in grapes, bud setting in aspen and more. Thus, instead of a designated flowering hormone florigen was established as a generic regulator of growth and termination (Fig. 1B and 1C, Shalit et al., 2009, Turnbull, 2011, Navaro et a., 2011, Lifschitz et al., 2014)

**Fig. 1.**
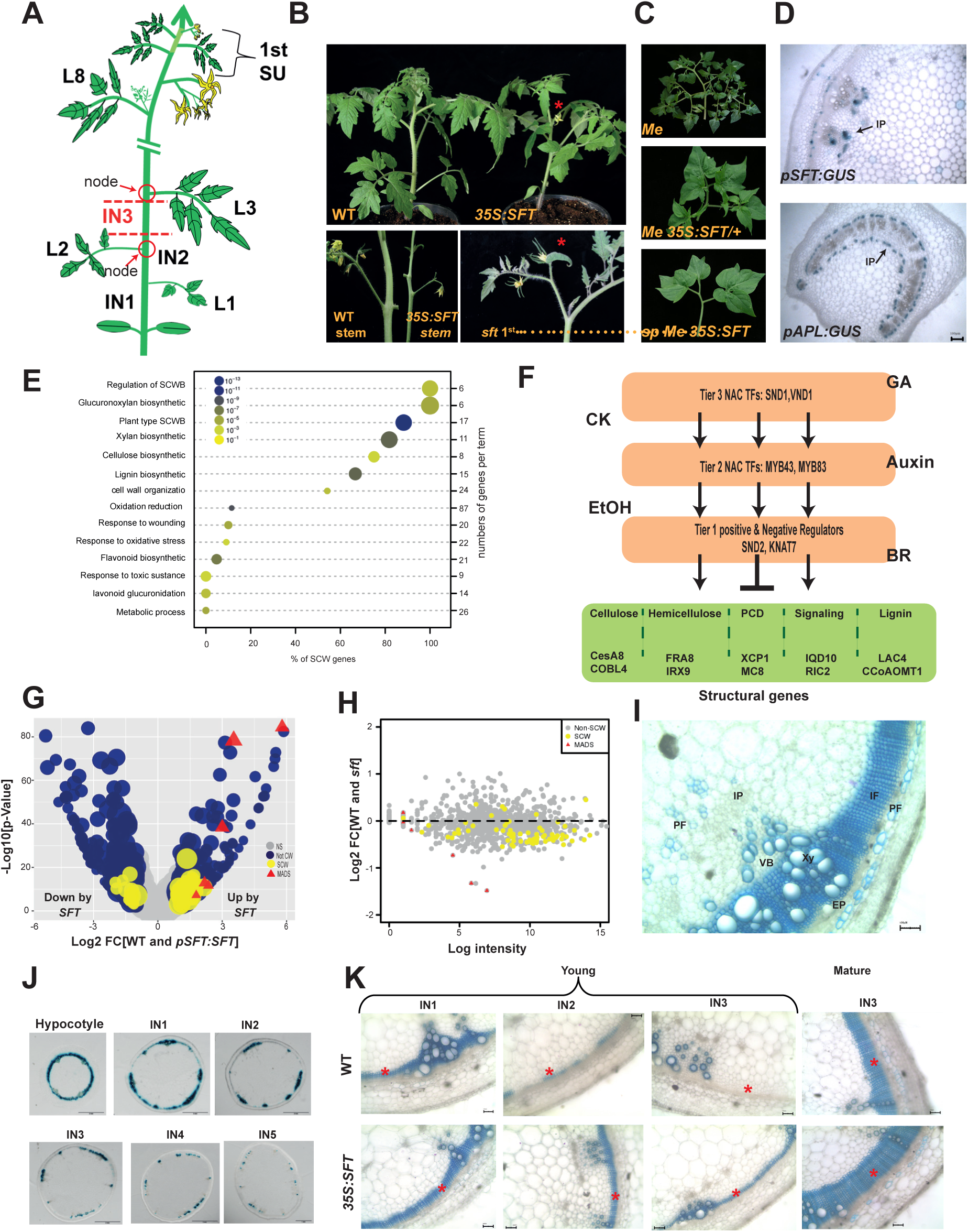
Florigen stimulates SCWB thereby restricting lateral expansion of tomato stems. (**A**) WT tomato: Leaf (L), Internode (IN), Sympodial Unit (SU). **(B**) *35S:SFT* induces premature flowering (Top) and slender stems (Bottom left). Bottom right – A single flower truss of *sft*, astriks indicate flowers **(C**) A gradual suppression of *Me* (*TKN2*) ramified compound leaves by increasing doses of florigen (*SFT*)**, (D**) Top-*pSFT:GUS* expression in external and internal stem phloem. Bottom-leaf petiole cross section showing p*TAPL:GUS* expression in both phloem layers. **(E**) Florigen regulates SCWB network: Functional classification of DEGs (padj<0.1 |FC≥2|) in *pSFT:SFT* vs. WT. Dot plot classifying top significant GO terms (M&M). **(F**) SCWB transcription network (after 21-24). **(G**) Florigen stimulates SCWB network and MADS TFs: Volcano plot comparing WT and *pSFT:SFT*. Circle size represents expression level. Grey-equally expressed. Color coded dots represent classes of 981 DEGs (padj≤0.1 |FC ≥ 2|). Blue - non-SCWB genes. Yellow - SCWB genes. Red triangles - MADS TFs. **(H**) SCWB network is down regulated in *sft* stems: distribution of 981 DEG (WT vs. *pSFT:SFT*) in *sft* vs. w.t, MADS and SCWB genes show a significant downregulation trend (p=8.1*10-14, t-test). **(I**) Principle vascular elements in a mature stem: VB-vascular bundle, Xy-xylem, PF-phloem fibres, EP-external phloem, IP internal phloem, IF-Interfascicular xylem fibres. TBO staining for lignin. **(J)** Tomato *pCESA4:GUS* along 5 week old stems. (See earlier stage in Suppl. Fig. 10). **(K)** Sections of 4 weeks, (young) and 40 days old (mature) WT, and *35S:SFT* - lignin staining. Asterisks indicate IF zones.

The transition to flowering marks a global, vegetative to reproductive, switch in the shoot system that is associated with altered growth homeostasis and modified shoot architecture, and the reallocation of resources and signals. However, the connection between phase reprograming and flowering and the genetic and molecular bases for the pleiotropic functions of florigen remain unknown. We hypothesized that if the mechanism for floral induction is universal, the cellular mechanisms underlying the global shift of the shoot system to reproduction would also be universal and thus navigated by the vegetative functions of florigen.

A search for a cellular mechanism underlying the pleiotropic morphogenetic effects of plant hormones was hindered for the lack of a common developmental denominator (Leyser, 2018). That all morphogenetic effects of florigen may be attributed to growth termination is thus unique. Unlike classic plant hormones, florigen performs identical developmental functions across species and has no metabolic intermediates or branching derivatives and is not essential for vegetative growth. The hormone is produced exclusively in the companion cells of the phloem (Zeevaart, 1976, An et al., 2004, Chen et al., 2018, Turnbull et al., 2013) and functions as a versatile molecular adaptor of transcription factors (TF’s, Pnueli et al., 2001, Taoka et al., 2011). Significantly, its abundance does not alter the designated fate of any of the organs involved, floral transition included. To execute a tunable switch from growth to termination and differentiation, florigen must regulate, non-disruptively, systems such as cell proliferation or cytoskeletal organization. Cellular systems, because transcription factors (TFs) are not the ultimate building blocks of cellular differentiation.

Our strategy for the unraveling of such systems consisted of initial deep RNA profiling designed to identify candidates for end cellular targets and then to trace the system back to the initial florigen signal. Here we focused on the ways florigen regulates the radial expansion of the stems. Stems consist of a limited number of cell types and their differentiating zones are entrenched in hierarchically demarcated developmental territories, circumventing the difficulties associated with the complexity of apical meristems (Fig. 1I, Taiz and Zeiger, 2010, Sparks et al., 2013). We discovered the acceleration of the SCWB transcription network and the consequential enhanced maturation of the conducing system as the mechanism used by florigen to coordinate the reprograming of the shoot with flowering. SCWB signifies a hierarchical transcriptional system (Fig.1F) that navigates and integrates the biosynthesis of the cellulose (Kumar and Turner, 2015) lignin and hemicellulose biopolymers (Hao and Mohnen, 2014) and guides their eventual deposition onto the inner layers of primary cell walls (Aloni, 2001, Caffey, 2002, Scarpella and Helariutta, 2010).

The mechanism underlying the arrest of lateral expansion of stems may well underlie the universal restriction of leaf growth under high florigen levels. Our data unveiled a target metabolic network co-opted by a plant hormone to regulate its fundamental function. This novel finding is accompanied by empirical evidence showing that flowering and SCWB are coordinately accelerated by florigen but are not the consequence of one another. The link between the flowering-inducing hormone and SCWB provides a novel paradigm of a global coordinating agent that is not involved in the regulation per se of any of the cellular systems involved.

## Results

### High SFT levels accelerate SCWB in the tomato stem

In tomato, high florigen levels stimulate precocious primary flowering while the pace of flowering during the sympodial phase is only marginally affected. By contrast, radial contraction of the stems is accelerated with age, in both the vegetative and reproductive phases, and is the single most robust pleiotropic effect of florigen (Fig. 1B, Lifschitz et al., 2006, Shalit et al., 2009). Further, stems are composed of a central vascular system comprised of a limited number of cell types, entrenched in hierarchically demarcated developmental territories surrounded by parenchyma cells thus circumventing the difficulties associated with the complexity of the SAMs (Patricka and Benfey, 2008, Taiz and Zeiger, 2010).

To unveil cellular systems targeted by florigen in stems we conducted an exploratory RNA profiling from the 3^rd^ internodes of 5-week-old WT., *sf*t and *pSFT:SFT* transgenic plants (Fig. 1A and 1B). Importantly, *SFT* is the sole contributor of florigen in cultivated tomato (Mc Garry et al., 2016). Comparison of the WT and *pSFT:SFT* transcriptomes (padj<0.1 and |DE>2|) revealed 981 differentially expressed genes (DEGs, Suppl. Table 1) significantly enriched for the metabolic and regulatory genes involved in SCWB (Fig. 1E and 1G). Consistent with this enrichment, 12 out of the 69 DE TFs encoded the tomato homologs of the *MYB* and *NAC* master regulators of SCWB (Hussay et al, 2013, Caffal and Mahonen 2009, Zhong and Ye, 2015). Surprisingly, 6 MADS genes, including four members of the *FUL* clan (Fig. 1G and Suppl. Table 2), known to mediate *FT-*dependent floral transition (Pin and Nilsson, 2012) were also upregulated in *pSFT:SFT* stems. Notably, other genes involved in flowering such as orthologs of *LEAFY* or *AGAMOUS* were not altered by florigen.

SCWB encompasses a hierarchical metabolic network that integrates the biosynthesis of cellulose lignin and hemicellulose and guides their deposition onto the inner layers of primary and secondary cell walls (Fig. 1F). If SCWB is a bona–fide target of florigen, SCWB-related transcripts should be underrepresented in *sft*. *TFUL–like* genes that were activated in *pSFT:SFT* stems, were indeed less expressed in *sft* (Suppl. Table 1). However, no significant enrichment of suppressed SCWB genes was detected in *sft* compared to WT. Nevertheless, the majority of SCWB genes upregulated by *pSFT:SFT* were expressed in *sft* stems at levels below that of the WT and this response was consistent and significant (Fig. 1H).

### Accelerated SCWB is correlated with precocious vascular maturation

To explore the anatomical manifestation of the enriched SCWB genes we followed secondary growth (SG) and deposition of SCW along WT and *SFT* overexpressing stems. The patterning of SCW deposition along tomato shoots follows the stereotypical basipetal gradient of eudicot plants (Hall and Ellis, 2012) as demonstrated here by the activity of the tomato p*CES A4* reporter (Fig. 1J). The bulk of the SCW is deposited during SG onto walls of the fascicular and interfascicular (IF) cells produced by the cambium which are fated to form the vessels and fibers of the secondary xylem and the primary phloem fibers (Fig. 1I, Aloni 2001,Chaffey et al., 2002). In practice we followed the formation and progressive lignification of the IF xylem fibers generated by the vascular cambium (Figure 1I). At 25 days post-germination (DPG), the first lignified IF xylem layers in the basal internodes of WT stems had just differentiated, while several lignified IF layers had already formed in the same internodes of the *SFT* overexpressing plants (Fig. 1K). A mature vascular system of *35S:SFT* stems is shown in Suppl. Fig.1C. These features correlated with the flowering status (Brukhin et al., 2003) of the tested genotypes: 9 leaves and stage 3 inflorescences in WT vs. 3 leaves and a fully blossomed inflorescence in *SFT* overexpressing plants. However, although vascular maturation correlates with flowering in WT plants it can proceed independently of flowering. The development of lignified xylem layers in the late flowering *sft* plants, and in the never-flowering *uf sft* double mutant genotype, were only marginally different from those from WT plants. Moreover, at 20 DPG, no lignified IF xylem rings were evident in the basal internodes of *SFTox* plants bearing stage 10 primary inflorescences (Suppl. Fig. 1C). The precocious activation of SG and SCWB, at both the molecular and histological levels, may fulfill the need for a comprehensive reallocation of signals and metabolic resources prescribed by the shift to the reproductive phase activated by florigen.

### Graft-transmissible florigen is sufficient to promote SCWB in recipient stems

The promotion of SCWB in the stems by high *SFT* reflects the combined impact of endogenous, stem-borne florigen (here after florigen) and of a mobile florigen (here after m-florigen) imported from the leaves. To identify cellular systems regulated by the m-florigen alone we studied five graft combinations, reporting the response of *sft* and *SFT* recipient genotypes to m-florigen contributed by WT and *35S:SFT* donors (Fig. 2A and 2B). The inherent variation associated with the graft assemblies, the cumulative and pleiotropic functions of florigen, the efficiency of mobility and the dynamics of vascular maturation (Hall and Ellis 2012, Brady et al., 2007, Taylor– Teeples et al., 2015) delineated low transcriptional responses and modest expectations. A group of 593 genes differing significantly between the WT and *sft* recipients of the homografts and the corresponding three heterografts were identified (Fig. 2C, Suppl. Table 3). Enriched GO terms for the 593 DEGs exclusively represented SCW functions (Suppl. Fig. 2A) and hierarchical clustering linked almost all DE SCWB genes into one cluster (Red in Fig. 3C). The clustering of the master regulators of SCWB with their target metabolic genes and that of the floral MADS genes with the SCW network depicted in Fig. 2C, strongly supported the hypothesis that m-florigen impacts SCWB as a network and that this role is mediated, at least in part, by the *TFUL* genes (Suppl. Table 2).

**Fig. 2.**
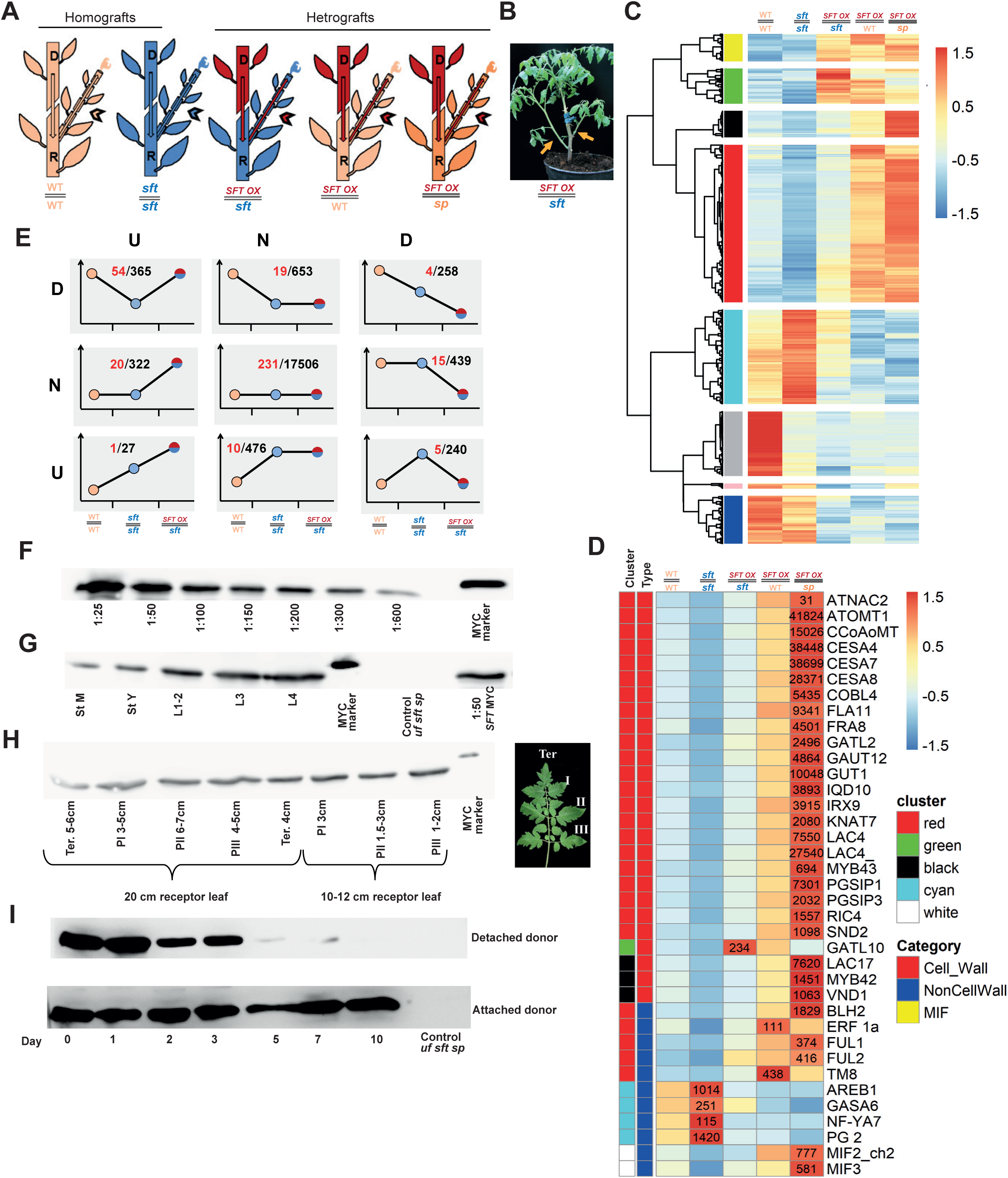
Graft-transmissible-florigen enhances the SCWB network in *sft* and WT recipients. (**A**) Grafts: Donor scions - upper and recipient shoots - lower part. Arrowheads – Sampled lateral recipient shoots. WT-light orange, *sft* – blue, *35S:SFT* red and *sp* is orange. (**B**) *35S:SFT//sft* graft.5 week old donors and recipients were used. Stems of lateral shoots (orange arrows) were harvested. (**C**) Hierarchical clustering of 593 m-florigen regulated genes. Most SCW genes are clustered in Red. (**D**) Heat-map of selected m-florigen-responsive genes from Figure 2C and their absolute expression values. SCW genes represent the major families in the network. Color coding is by cluster as in Figure 2C and by CW, non CW and MIFs. (**E**) Punnett Group (PG) - Expression patterns of the left three grafts defined by two bipartite comparisons WT//WT vs *sft//sft* and *SFT//sft* vs *sft//sft* and categorized: Up (U), Down (D) and No change (N), |FC≥1.5| (M&M). Cohort DU (upper left) is enriched for the florigen–regulated genes analyzed in Fig 2C (101/593, p< 5.4*10-70 hypergeometric) and for SCWB genes (54/ 365, p< 4.4*10-26). Circle color coding corresponds to recipient genotypes in 2A. Red numbers: SCWB genes. (**F**) A dilutions series of *SFT*-3XMYC from donor *35S:SFT-*3XMYC leaves. (**G**) m-florigen accumulates in recipient leaves and stems at about 1/100 of donor leaf levels. L1- old leaves. St M and St Y, mature and young stems respectively. (**H**) MYC-tagged m-florigen is distributed among all leaflets of mature recipient leaves. P –pair of leaflets. Ter - terminal leaflet, most mature (see on the right).(**I**) m-florigen has a ca. 4 day half-life. Top: Daily recovery of the mobile *SFT*-MYC in recipient leaves after donor removal. Bottom: Recovery of the mobile *SFT*-3xMYC from recipient shoots of intact grafts.

**Fig. 3.**
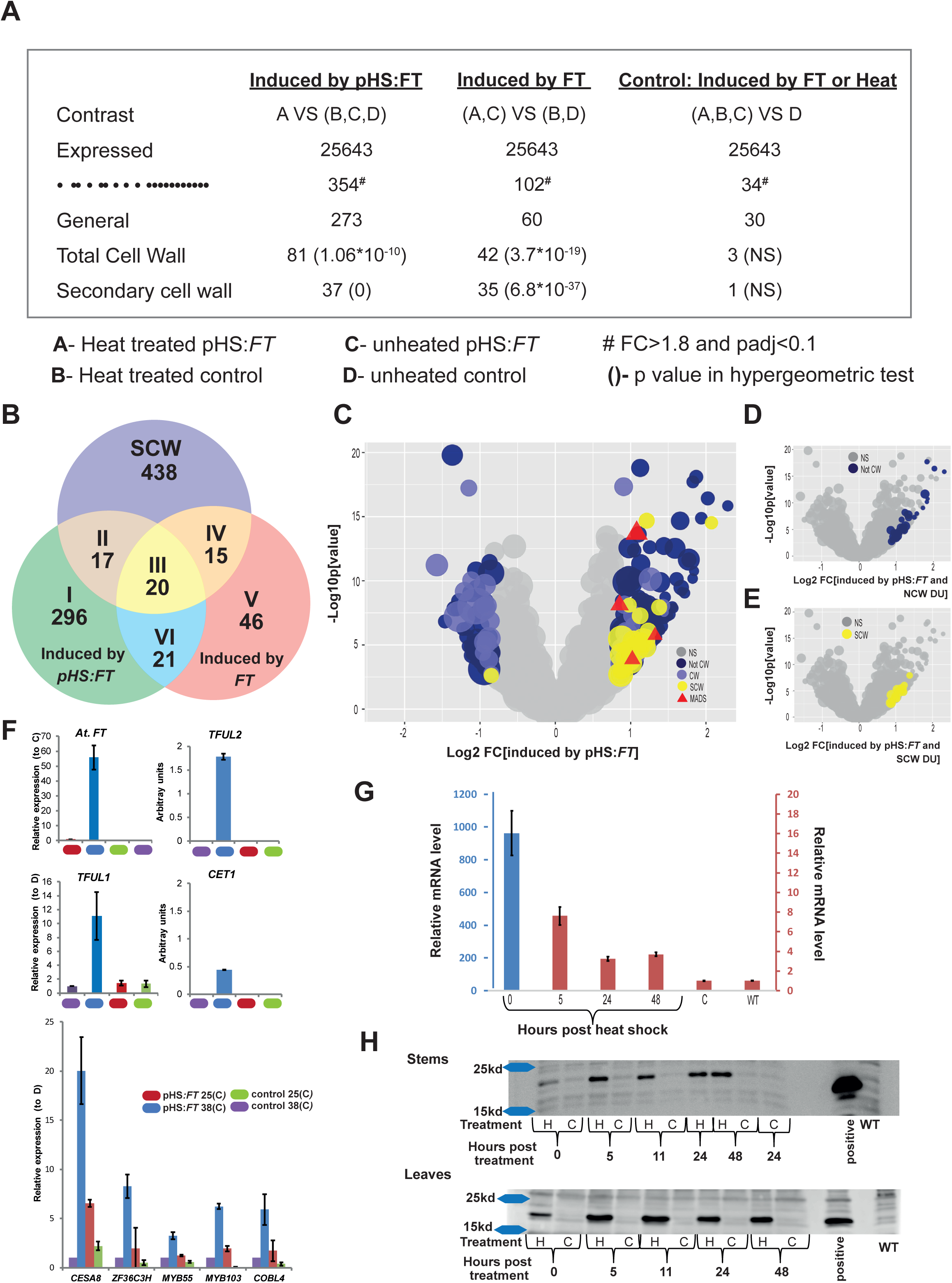
Transient heat-shock induction of *AtFT* significantly stimulated SCWB genes in tomato stem. (**A**) Identification of heat-shock (HS) induced *FT*-responsive genes. One factor design comparing RNA profiles of heat treated *pHS:FT* stems vs the combined control treatments with padj<0.1 and |FC>1.8| (A vs B,C and D). Comparisons to find *FT* leaky regulated genes (A and C, vs B and D) and heat induced genes (ABC vs D) were also made. **(B**) Venn diagram showing significant overlap of *FT* - induced genes with SCW genes in tomato. **(C**) Volcano plot comparing DEG in induced *pHS:FT* and all other control conditions. (padj≤0.1, |FC≥ 1.8|). (D-E) *FT*- induced genes overlap with 55 non-CW (D) and 23 SCW (E) genes from cohort DU of PG (Fig 2E), also up-regulated.**(F**) qRT-PCR validation of 9 genes activated by heat-induced *AtFT* in tomato stems. (G) Time course - activation and degradation of HS-induced *FT* RNA in tomato stems: The initial two order of magnitude higher expression level of *FT* mRNA is at the 3-5 fold level after 5 hours. (H) HS-induced SFT-FLAG protein persists for at least 48 h in stems (top) and leaves (bottom).

Consistent with m-florigen functioning primarily as a booster of a functioning SCWB system, quite similar to its impact on flowering, SCW and non-CW genes regulated by florigen showed preferential response in the *SFT* recipients (grafts 4 and 5) as compared to the *sft* recipients (Fig. 2A). Thus, while moving along the phloem track, florigen emits signals that enhance the SCW metabolic network in the stems.

The observed enrichment for SCWB genes hinges on their functional classification. In an alternative analytical approach we performed tripartite comparisons between recipient genotypes: Here, all 20,286 expressed genes were ranked solely by their expression trajectories and then divided into nine categories visualized in what we refer to as Punnett Grids (PG). In the most instructive tripartite comparison (PG, Fig. 2E) we compared the response to m-florigen of *sft* recipients in the two bipartite contrasts WT//WT vs. *sft*//*sft* and *35S:SFT*//*sft* vs. *sft*//*sft* (See legends and M&M). Cohort DU of PG (upper left) includes 365 genes (Suppl. Table 4) that were down regulated (<u>D</u>) in the recipient *sft* stems in the first bipartite contrast but which were “rescued” (Up-U) by m-florigen in the recipient *sft* stems of the second bipartite contrast. Functional annotation of these 365 genes revealed an exclusive enrichment of SCWB genes (Suppl. Fig. 2B). At this stage we assembled a curated list of 498 SCW tomato genes, representing 265 Arabidopsis genes (M&M), and tested their distribution across the 9 cohorts of PG (Red numbers in PG). Of the SCW genes expressed in the graft experiments 54 (hyper geometric p-value 5.9×10^−33^) were assigned to cohort DU with no preferential presentation in any other cohort (Fig. 2E). Thus, many genes responding to florigen in *sft* and *SFT* backgrounds represent SCW functions. SCWB emerged as the most likely metabolic target of **m**-florigen in tomato stems. Consistent with florigen functions as an accelerator of an already operating network, genes involved in the specification or patterning of SCWB (Cano Delgado et al., 2010, Scarpella and Helariutta, 2010) were not regulated by high florigen.

### The mobile florigen is extremely stable, moves un-modified and invades all aerial organs

In prevailing descriptions, the vascular system is conceived merely as a conduit for florigen on its rectilinear passage to the apical meristems. Here we showed that the stem vasculature, particularly the xylem, is also a target for the hormone (Fig.1). We thus studied the molecular constitution, organ distribution and stability of m-florigen. Mass-spectrometry analysis suggested that affinity-enriched SFT-MYC from recipient leaves (M&M) moves free of major post–translation modification. In contrast, the antagonistic SP-MYC protein was phosphorylated in Serine 157 which is lacking in SFT (Suppl. Fig. 2C and 2B).

By 25 days post-grafting, the MYC-tagged transmissible florigen was detected in the stems and in every single leaflet of mature and young leaves of the recipient, at approximately 1/100 of its level in the donor leaves (Fig. 2G, H). Note that a single donor leaf is sufficient to promote lasting flowering in *sft* recipient plants (Lifschitz et al, 2006) and that the expression level of *SFT* in *35S:SFT* plants never exceeds 1/20 th of the promoter potential (i.e. ca. 1000 vs. 25000 DESeqs for other genes). A rough estimation of the quantities of the transmissible mobile protein yielded, depending on the tissue, 14-140 ng per 1gr of initial tissue which seems quite high considering the low expression level in the donor. This, together with the ‘self-propagation’ phenomenon of florigenic signals (Zeevaart, 1976) and the quantitative nature of floral stimulation, led us to examine the stability of the mobile hormone. Indeed, upon removing donor scions from *35S:SFT-MYC//sft* assemblies 21 days post grafting and monitoring SFT-MYC in recipient leaves, florigen was shown to have an exceptionally long half-life of approximately 4 days! (Fig. 2I, and see Kim et al., 2016). The long range mobility, extended stability and indiscriminative distribution of the protein among all vegetative organs are consistent with an auto–regulatory model for the allocation and abundance of florigen (Shalit et al., 2009).

### Transient heat induction of the Arabidopsis *FT* gene stimulates SCW genes in tomato

Our hypothesis that florigen guides the dynamics of SCWB is contingent on steady state transcription profiles. To determine whether florigen impacts SCWB on a shorter time scale we used the *pHS:FT* transgene (*pHEAT SHOCK:FT*, a gift of O. Nilsson) to create *sft pHS:FT* plants bearing a heat-inducible florigen.

The experimental scheme was comprised of heat treated *sft pHS:FT* plants (A) and 3 control treatments: untreated *sft pHS:FT* (B), heat-treated *sft* (C) and (D), untreated *sft* plants (Fig. 3 A). Five week old plants were exposed for 90 min to 38° C hot air and stem segments from their 3^rd^ internodes were harvested 24 h later (M&M) to allow for the recovery of normal transcription and accumulation of florigen. Of the 354 DEGs identified 24h after heat treatment (Fig. 3A), 37 were SCW-related, 44 were CW-related and 273 were non-CW genes including *TFUL1* and *TFUL2* (Suppl. Table 5). Genes induced by heat treatment or by basal expression in *pHS:FT* plants significantly overlapped with the list of tomato SCW genes (Fig. 3B) and the *FT*-induced SCW genes as well as 2 MADS genes were all upregulated (Volcano plot in 3C). qRT-PCR validation of 9 FT-induced SCW genes is shown in Figure 3F. Comparison of the 273 *pHS:FT*- activated genes and the 365 m-florigen-regulated genes of cohort DU of PG (Fig. 2E) revealed that 23 of the 37 SCW and 55 of the 273 total genes activated by *FT* were part of the DU cohort of PG. Some of these 55 genes may represent functions not previously associated with SCW or florigen.

To investigate the fate of heat-induced FT transcripts and polypeptides we created *pHS:SFT-FLAG* plants. More than 95% of the *SFT-FLAG* transcripts presented at the end of the 90 min heat treatment vanished 5 h later. The remaining transcripts were approximately 3-fold higher than control levels at 48 hours post-treatment (Fig. 3 G). In contrast, 5h following a 90 min heat treatment the level of the SFT-FLAG protein was two-fold higher in both stems and leaves, for at least 48 h (Fig. 3H). Thus, florigen, in its endogenous or mobile forms, can trigger the upregulation of the SCWB network in tomato stem before any anatomical changes are evident- as early as 24 hours post *FT* induction. The activation of SCW genes in the day-neutral tomato by *FT* from photoperiodic Arabidopsis suggests that boosting SCWB, like flowering, is a universal property of florigen.

### The *Ginkgo biloba FT*: A mobile antagonist of flowering and a suppressor of SCWB provides a molecular portrait of an anti-florigenic syndrome

In its role as a floral enhancer, florigen is checked by systemic antagonistic systems (Zeevaart 1976, Lifschitz et al., 2014). If florigen deploys similar mechanisms in boosting SCWB in stems, it is expected that its floral antagonists will also suppress SCWB. Several years ago, out of curiosity, we created transgenic tomato lines overexpressing *FT*-like genes from the gymnosperms *Ginkgo biloba (GinFT)*, *Cycas rumphii* (*CrFT*) and *Piscea sitchensis*. Confirming reports on floral suppression by gymnosperms *FT-* like genes (Klintenas et al., 2012, Liu et al., 2016), *GinFT* generated the most extreme anti-florigenic syndrome in tomato, Arabidopsis, and tobacco (Fig. 4A-D and Suppl. Fig.4). In addition, *GinFT* plants feature shorter internodes, complementation of *sp*, leafy inflorescences and leafy sepals. GinFT, like SP, interacts with the Arabidopsis FD and its tomato homolog SPGB (Suppl. Fig. 4E, F, Pnueli et al, 2001). Yet, the most pertinent hallmarks of the *GinFT* syndrome were the extreme age-dependent radial swelling and twisting of the stems and leaf petioles (Fig. 4A-C).

**Fig. 4.**
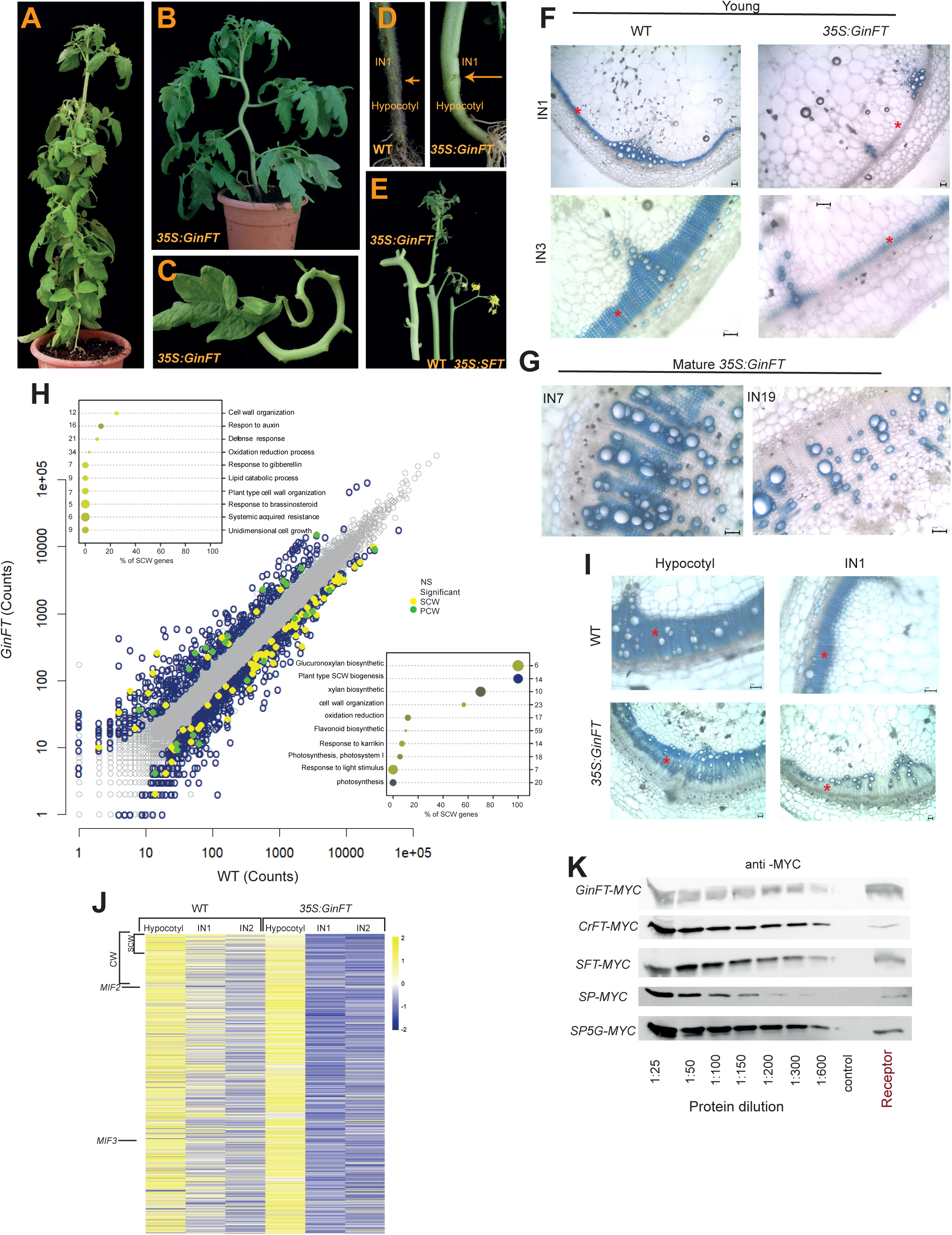
*GinFT* antagonizes florigen, suppresses SCWB and prolongs vascular maturation. (**A**) Adult *35S:GinFT-MYC(GinFT)* transgenic plant with condensed leaves. **(B**) Young, a 4- week-old *GinFT*plant with a convoluted stem but early expanding leaves. **(C**) The petiole and rachis of a mature curled *GinFT* leaf. (**D**) Hypocotyls of *GinFT* plants undergo differential radial constriction. Arrows – hypocotyl-stem junctions. **(E**) Radial expansion of inflorescence-bearing stems in G*inFT*, WT and *35S:SFT* plants. **(F-G**) High expression levels of *GinFT* delay lignification of IF fibers. F) Top - 30 day old WT and *GinFT* plants. Bottom - 40 day-old WT and *GinFT* plants. **G**) Excessive non-lignified IF xylem fibers of 70 day-old *GinFT* plants. **(H**) SCWB genes are significantly downregulated in *GinFT* stems. Scatter plot displaying expression levels of *GinFT* vs WT. Colored circles represent DEGs (padj<0.1, |FC>2|). Insets: Top 10 GO terms in upregulated (top-left) and downregulated (bottom right) DEGs. **(I**) SCW deposition, hypocotyls and stems exhibit differential sensitivity to high *GinFT* expression. Note the accumulation of non-lignified IF xylem fibres in *GinFT.* Cross sections, TBO staining. **(J**) Heatmap of 1698 DE genes (FC≥2) between the hypocotyls and internode 1 from *35S:GinFT* plants shows unique developmental differences in growth regulated by *CETS*. **(K**) Gymnosperm CETS proteins are mobile. MYC-tagged protein dilution series from the indicated donor tissues and the undiluted mobile CETS proteins from the corresponding recipient shoots.

To understand these phenotypes we performed histological comparisons of SCW deposition in both stems and hypocotyls. Unlike WT plants (Fig. 4F), the basal internodes of 30 day old *GinFT* stems were devoid of concentric lignified IF fibers. At 75 DPG, while fully blossoming, lignification of secondary fibers in the basal internodes of the stem were still significantly delayed (Fig. 4G).

To see if *GinFT* suppresses SCWB as a network, we profiled RNA from the 1^st^ (Bottom) internodes of 35 day old WT, and *35S:GinFT* plants. GO analysis of the 1122 DEGs, (representing 976 A.t. genes, FC≥2) revealed a significant enrichment of SCW genes. This time however, the majority of SCWB genes were downregulated rather than upregulated, providing a molecular portrait of an anti-florigenic syndrome (Fig. 4H, Suppl. Table 6).

Interestingly, in addition to the swelling of stems we noticed a differential radial contraction of the hypocotyls of *GinFT* plants (Fig. 4D), which, reflects a proportionally enhanced differentiation of cambial derivatives in hypocotyls of *GinFT* plants (Fig. 4I). Consistent with the histological phenotype, the slight global expression gape between hypocotyls and stems in WT plants is biased in favour of SCW genes in the hypocotyls of *GinFT* plants (Fig. 4J).

The evolutionary choice of *CETS* genes for the systemic induction of flowering and the antagonistic functions of *GinFT* in tomato implied that such genes were already endowed with pertinent auxiliary potentials in non-flowering plants. We thus hypothesized that *CETS* genes may already co-ordinate vascular maturation, perhaps long-range, in gymnosperms. Consistent with this speculation, the MYC-tagged polypeptides produced by *35S:GinFT-MYC* and *35:CrFT-MYC* transgenic plants were as mobile as tomato CETS proteins encoded by the *SFT*, *SP* or *SP5G* genes (Fig. 4K). It is tempting to speculate that the link between CETS genes and SCWB underlies the annual growth cycles in the trunks of perennial trees.

### *TFUL2* functions downstream of florigen to regulate the dynamics of SCWB independently of flowering

During floral transition, florigen forms a regulatory network with MADS factors of the *FUL* clade (Fornara et al., 2009). If florigen exploits an analogous downstream operation when boosting stem SCWB, then *TFUL2*, which consistently responds to all forms of florigen, might be expected to accelerate SCWB in stems and in a florigen- and *SFT*-independent manner. A role for *FUL-like* genes in SCW processes was previously reported in fruits from both Arabidopsis and tomato (Ferrandiz et al., 2000, Fujisawa et al., 2014, Wang et al., 2014). Notably we found that 3*5S:TFUL2* plants developed extremely slender shoots and flowers after 5-6 leaves. WT progenitor plants flowered after 9 leaves, *sft 35S:TFUL2* plants after 11 leaves and *sft* plants after 12-13 leaves (Fig. 5 A-C). Four weeks post germination, 2-3 three-layer-thick, lignified IF fibres populated the two basal internodes of WT and *sft* plants but 10-11 layers were already produced in the same internodes of *35S:TFUL2* and *sft 35S:TFUL2* plants. Remarkably, the late flowering *sft 35S:TFUL2* stems were nearly as slender as those from the *35S:TFUL2* plants and their vascular maturation was similarly accelerated (Fig. 5D).

**Fig. 5.**
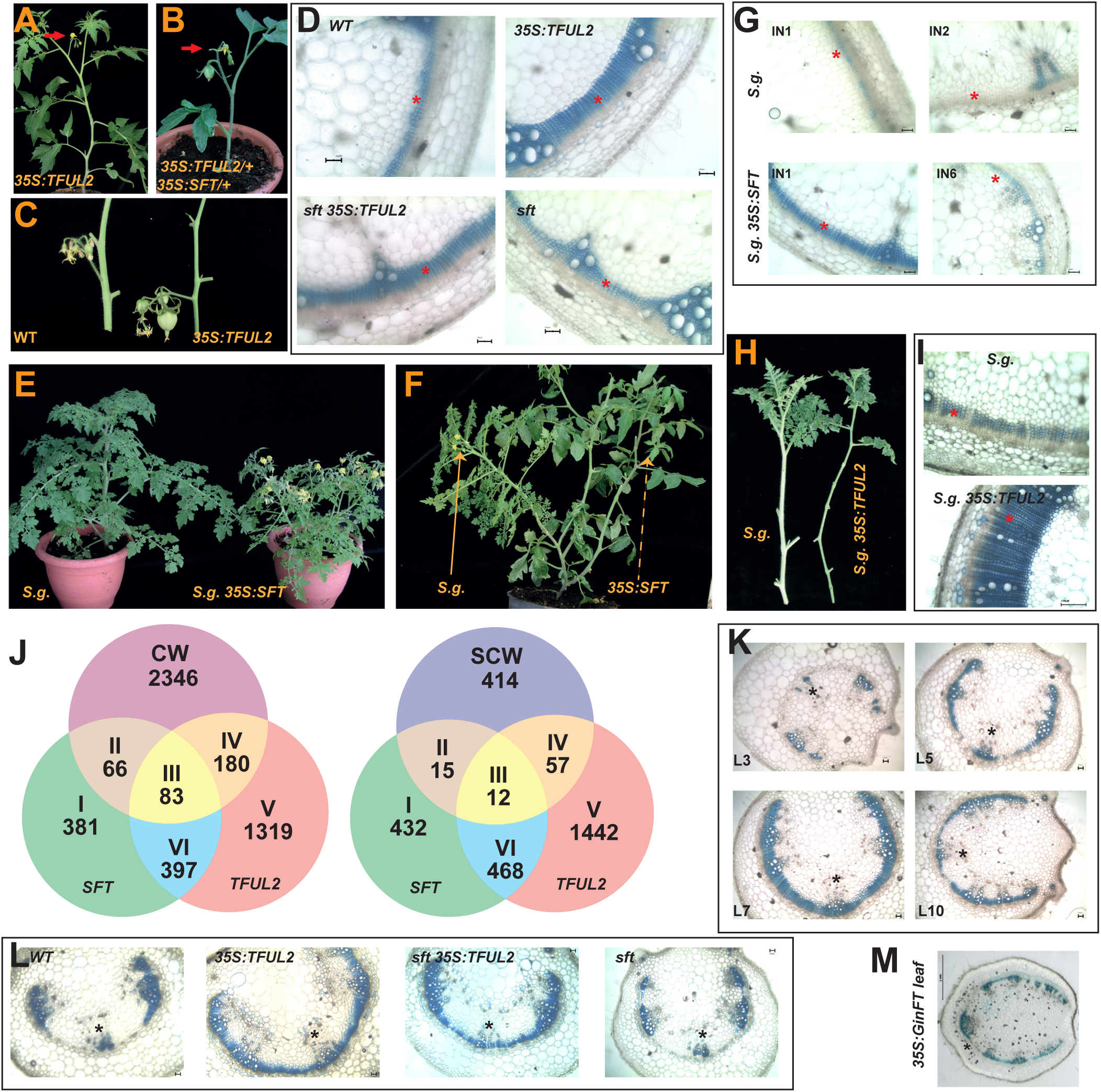
*TFUL2* induces premature, florigen-independent, SCW deposition. (**A**) Primary flowering (5 leaves) in *35S: TFUL2* vs. 2-3 in *35S:SFT* and 8-9 in WT. **(B**) Additive effects of *TFUL* and *SFT* on stem size and flowering. **(C**) Slender stems and beaked fruits in *35S:TFUL2*. See also Suppl. Fig 5H. (**D**) *TFUL2* induces flowering-independent acceleration of secondary growth and SCW deposition. Cross section of the first internodes of 30 days old WT, *sft* and *35S: TFUL2* plants. **(E**) *35S: SFT* induces flowering under long-days in 6 weeks old *Solanum galapagense* (*S. g*.). **(F)** Graft-transmissible florigen stimulates flowering in *S.g* under long days. **(G)** *35S:SFT* induces precocious SCW deposition and vascular maturation in S. *g*. grown under long-days. Compare internode 6 of *35S:SFT* with internodes 1 and 2 of WT *S.g* plants. **(H)** Shoots of non-flowering WT and *35S:TFUL2 S.g* grown under long days. **(I)** Cross-sections from WT and *35S:TFUL2 S.g* stems shown in Figure H. The external phloem fibres (PF) in *S. g. 35S:TFUL2* are already lignified. **J)** Venn diagrams showing overlapping of DEGs of *35S:SFT* or *35S:TFUL2* with CW (Left) or SCW (Right) genes in internode 1 stems. **K**) The distinct maturation gradient of petioles along the shoot. Note the absence of a complete xylem crescent in the most mature leaves. Asterisks mark mid-veins. **(L)** Cross sections of petioles from leaf 4 of 5 week old young plants of four genotypes stained for lignin. **(M)** *GinFT* leaf petiole of 80–day old plant.

Tomato is a day-neutral plant. To determine whether the florigen-SCWB link is valid in photoperiodic plants we created *Solanum galapagense* (*S.g.*) (a short-day wild relative of tomato), overexpressing *SFT* or *TFUL2* plants. We validated that overexpression of *SFT*, as well as a grafted *35S:SFT* donor, induce extensive flowering in WT *S. g.* under long days (Fig. 5E and 5F). Under long-days, WT *S.g.* showed a protracted delay in vascular maturation but *S. g. 35S:SFT* plants developed slender stems and enhanced xylem differentiation (Fig. 5G). By the same token, two-month-old *S.g.35S:TFUL2* plants, grown under long days and lacking signs of floral primordia, formed extreme slender stems (Fig. 5H) with fully differentiated IF xylem fibres (Fig. 5I). Taken together, the florigen- *TFUL2* module is valid in stems of a photoperiodic species and the rate of secondary xylem differentiation is correlated primarily with the expression levels of *TFUL2* and not with flowering per se.

### *SFT* and *TFUL2* regulate common and gene-specific cellular programs

Since high florigen levels activate *TFUL2* in stems, we studied the differences and commonalities in the ways *TFUL2* and *SFT* affect vascular maturation. To this end, we compared the transcription profile of their 1^st^ (bottom) internodes with that of WT plants. With a ≥2-fold change in relation to the WT, we identified DEGs for each overexpressing genotype and then calculated the overlap of genes regulated by *SFT* and *TFUL2*. We then superimposed them with our reference lists of CW and SCW genes (Venn diagrams in Fig. 5J, Suppl. Table 7). *SFT* was upregulated 3-fold in *35S:TFUL2,* suggestive of a positive feedback loop, as part of a primary module for the acceleration of SCWB. A significant number of CW and non-CW genes were differentially regulated by either *SFT* or *TFUL2*. The GO analysis of all classes of DE genes included in diagram 5J, indicated that the lignin metabolism responded differentially to *TFUL2* (Suppl. Fig.5 B-F). Interestingly, 80 of the 83 CW genes regulated by both *SFT* and *TFUL2* (class III, left diagram), responded in the same direction. Non-CW genes regulated by *SFT* or *TFUL2 (*class V, VI and I) may represent auxiliary activities consequential to the acceleration of SCWB, as well as functions previously not associated with SCWB.

If, as we surmise, florigen accelerates SCWB, it is expected to do so throughout the entire plant, including leaves. In tomato, the vascular strands of the petioles, the leaf stalks, undergo extensive SG, generating, at maturity, a lignified crescent-shaped IF xylem band. Surprisingly, unlike the basipetal maturation pattern along the stems, that of petioles displayed a bell shape configuration, which peaked in leaf 7 of 85-day-old WT plants (Fig. 5K). The 3-4 basal leaves of WT plants never form complete xylem crescents, although their corresponding stem internodes are the first to mature. Consistent with the acceleration of vascular maturation in *TFUL2* overexpressing stems, leaf 3 and 4 of 45 days old *sft 35S:TFUL2* and *35S:TFUL2* successfully completed entire xylem crescents (Fig. 5L). Conversely, not a single leaf in the 100-day-old *35S:GinFT* plants formed a continuous xylem crescent (Fig. 5M).

### The *MIF3* gene mediates vascular maturation downstream and independent of florigen

RNAseq data revealed SCWB as a target for florigen. To be valid, other genes not previously known to be associated with SCWB, but inferred from the same data set to be regulated by florigen, are expected to be involved in SCWB. To test this supposition we CRISPR-edited (Soyk et al., 2017) the two *MIF* (mini zinc fingers) genes (Hu and Ma, 2006, Sicard et al., 2008) that were activated by *SFT* and *TFUL2* but practically silenced, in *GinFT* stems (Fig. 6A). CRISPR-cas edited *mif2^cr1^* and *mif3^cr1^* (Suppl. Fig.6 and M&M) induced swollen stems and petioles (Fig. 6B) and delayed lignification of secondary xylem fibres in stems and leaves (Fig. 6C, 6D and 6E). These defects were reminiscent of *GinFT* plants but with two important differences: Firstly, the overall stature, flowering time and productivity in *mif2^cr1^* and *mif3^cr1^*plants were essentially normal. Secondly, LOF *MIF* genes conferred a differential tuning of SCW deposition in hypocotyls and stems similar to *GinFT* (Fig. 6F), but not a disproportional contraction of the hypocotyls. While *GinFT* promotes a direct developmental antagonism of the SFT-TFUL module, the *MIF* genes may regulate an intermediate sub–group of SCWB.

**Fig. 6.**
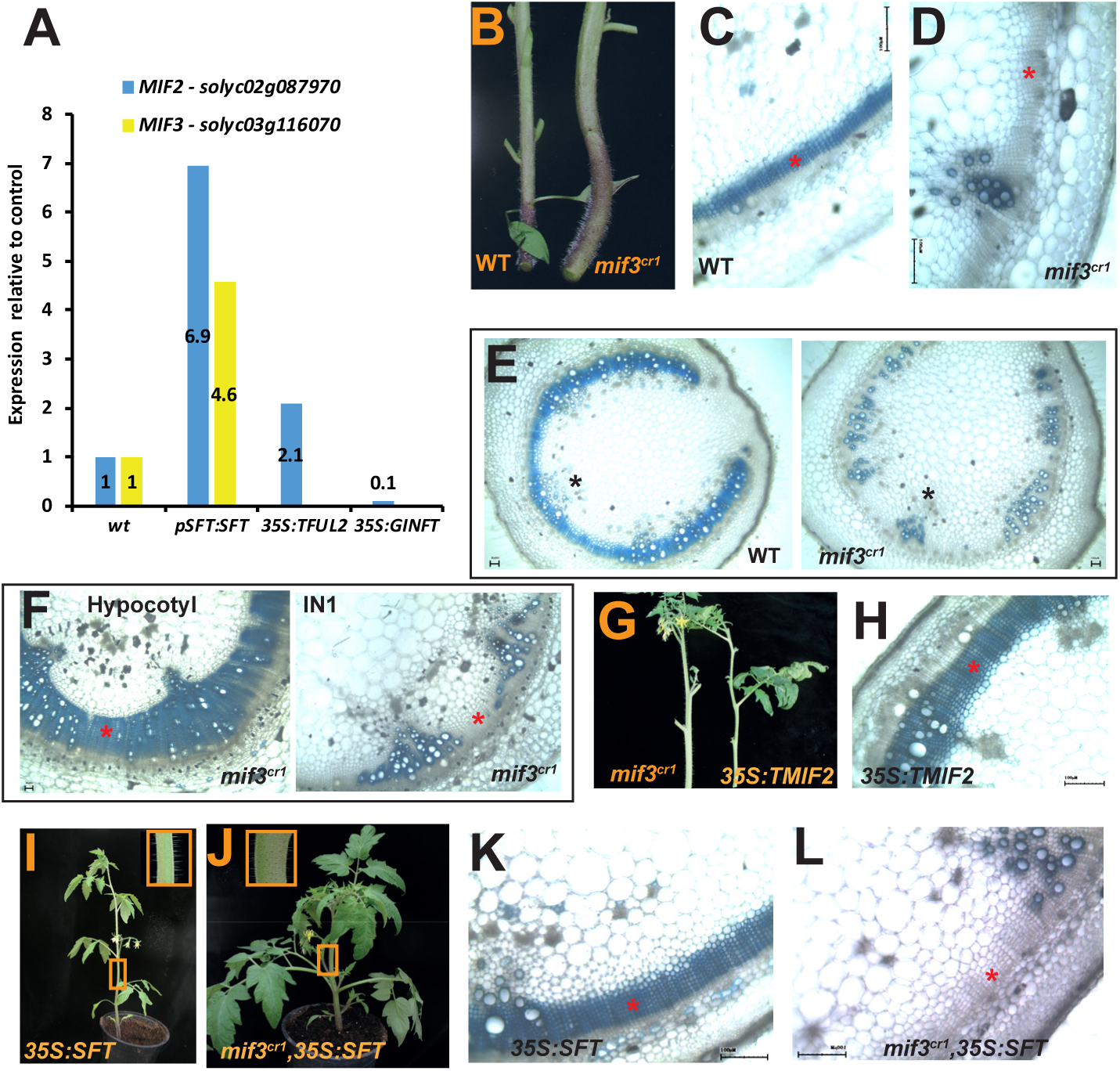
*MIF2* and *MIF3* relay florigenic signals to regulate SCWB. (**A**) Regulation of Tomato *MIF2* and *MIF3* by *SFT, TFUL2 and GinFT*. **B-D**) Radial expansion (B), SG and lignification in WT and *mif3^cr1^* stems.**E**) Xylem crescents in petiole No. 7 of 7 week old WT and *mif3 ^cr1^* stems. (**F**)A biased hypocotyl/stem differentiation in *mif3^cr1^*. For normal ratio consult Fig.4I.(**G-H**) Slender stems (G) precocious vascular differentiation (H) in *35S:MIF2-FLAG* plants. **(I-J)** Early flowering but thick stems in *mif3 cr1 35S:SFT* plants. (**K-L**) *mif3 ^cr1^* suppresses early SG and SCW deposition in *35:SFT* plants. See also Suppl. Fig. 6E.

Overexpression of *MIF* genes in Tomato and Arabidopsis induce growth retardation and malformed reproductive organs. Fittingly, the regulatory regions of the *MIF* genes in all three species contain putative MADS binding sites (Sicard et al., 2008). To compare the loss and gain-of functions in stems, we prepared *35S:MIF2–FLAG* and *35S:MIF3* plants and found that both genes induced rigid slender stems with an extensive, precociously lignified, secondary xylem (Fig. 6G and 6H) and a normal flowering time in WT and similarly in a *mif2^cr^ mif 3^cr1^* background.

The *MIF3* LOF alleles were generated in the *SFT* background. To determine the effect of higher doses of florigen we bred *mif3^cr1^ 35S:SFT* plants. Like *35S:SFT* plants, 32 days old *mif3^cr1^ 35S:SFT* flowered after 3 leaves but their stems remained as swollen as in *mif3^cr1^* plants (Fig. 6I and 6J). Furthermore, lignification of the IF cambial cells were similarly delayed and the hypocotyl-stem ratio was as wide as in *mif3^cr1^* alone (Fig. 6 K, 6L and Suppl. Fig.6). Therefore SCW deposition and xylem differentiation are independent of flowering time and flowering is uncoupled from vascular differentiation. In regulating SCWB downstream, and developmentally independently, of florigen, the two *MIF* genes represent a new regulatory tier partially bridging the gaps between the florigen and SCWB in tomato stems.

## Discussion

Floral induction, in addition to creating reproductive organs, transforms the shoot system from the vegetative to the reproductive phase. The reproductive phase entails reprograming of metabolic programs, moderation of vegetative growth and amendment of the global source - sink relations. We hypothesized that if the mechanism for floral induction is universal, the mechanisms navigating the shift to reproduction would also be universal and that florigen itself may activate it. We show that concomitantly with the induction of flowering florigen independently accelerates vascular maturation in stems and leaves. The developmental choice to reprogram global growth by coordinating floral induction with vascular maturation is logical. The continuous growth of plant organs hinges on cell divisions and cell expansion: both contingent on the composition of the cell walls (Cosgrove, 2005, Yang et al., 2016). Enhanced mechanical support, more rigid vasculature and efficient water conductance meet the emerging needs of the reproductive phase. Furthermore, precocious vascular maturation will inevitably promote global reprograming of the shoot system and the consequential redistribution of resources and signals in the adaptation to changes enforced by the transition to flowering. Unveiling the unanticipated link between florigen and vascular maturation was experimentally rooted in the genetics of tomato, the low complexity of the stems, the resolving power of RNA sequencing and, critically, the elucidation of the transcription network navigating SCWB (Figure 1F). The enrichment for SCWB genes in four independent experimental platforms, and the significant overlap of the cohorts of genes enriched in each experiment, substantiate the hypothesis that florigen enhances SCWB through a gene regulatory network. Furthermore, florigen, in its endogenous or mobile forms, can trigger the upregulation of the SCWB network in tomato stem before any anatomical changes are evident- as early as 24 hours post *FT* induction (Fig 3). Consistent with florigen functions as an accelerator of an already operating network, genes involved in the specification or patterning of SCWB (Caño-Delgado et al., 2010, Lukas et al., 2013) were not regulated by high florigen. To accelerate SCWB florigen activates MADS factors of the *AP1/FUL* clade it employs universally in boosting flowering (Fornara et al., 2010). The ability of *TFUL2* to enhance SCWB in the *sft* background was the first indication for the independent role of florigen in regulating these two pivotal processes. This prediction was then supported by the relation between flowering and vascular maturation rate in the short day *Solanum galapagense*. Moreover, overexpression of *GinFT* and inactivation of *MIF3* had similar suppressing effects on vascular maturation but contrasting effects on flowering time. Functional *FT-like* genes are already present in gymnosperms (Liu et al., 2016). The evolutionary choice of a *CETS* gene for the systemic induction of flowering implies that such genes were already endowed with pertinent auxiliary potentials in non-flowering plants. In agreement, *GinFT* displays systemic potential, delayed floral transition and attenuated SCWB, suggesting that a long-range regulation of vascular development in gymnosperms by *CETS* genes offers such an advantage. It is interesting to speculate that the link between *CETS* genes and SCWB lies in the annual growth cycles in the trunks of perennial trees. Several observations suggest that Florigen impacts shoot architecture by regulating growth and termination in every aerial meristem but that it is not essential for any morphogenetic function: A) The SAM grows normally without florigen and transitions to flowering via florigen. B) *sft* tomato and *ft, tsf* Arabidopsis plants eventually flower without florigen (Yamaguchi et al., 2005). C) Florigen is not essential for normal vascular development but vascular maturation is enhanced by it and each organ, leaf, stem, and hypocotyl displays an organ-specific response to *SFT/TFUL*, *GinFT* or *mif* genes (Figs. 4, 5 and 6). D) Compound leaves develop normally without florigen. It has remained overlooked however that, of all the potent transcription factors (Hay and Tsiantis 2010), a change in the proportion of florigen to antagonist (i.e. an elevated *SFT/SP* ratio) is the most effective suppressor of leaf complexity (Figure 1C). The florigen-SCWB link may thus offer an additional regulatory tier for systems such as leaf size and shape.

The realization that florigen independently boosts flowering and accelerates vascular maturation provoked “harmonization” as the term best representing its regulatory role. Harmonization refers to a coordinated progression of global developmental processes which otherwise proceed independently but, even when synchronized, are not the consequence of one another. Harmonizing agents function as “peripheral” regulators-peripheral to the basic tenets of the plants, not in reference to spatial operation or importance. The systemic florigen functions as a ‘peripheral’ regulator of the reproductive phase.

## Materials and Methods

### Plant material

Plants were grown in a glasshouse under natural conditions. The WT cultivars NY, M82 (*sp* background) and Money Maker (MM) were obtained from the C. M. Rick Tomato Genetics Resource Center at UC Davis. Other lines mentioned in the text were bred for this work. Transgenic plants were generated as described (Mc Cormick, 1991).

### Cloning procedures

*SFT* was amplified from cDNA with the appropriate primers containing EcoRI at the 5’ and SmaI at the 3’ end (as listed in Suppl. Table 9) and cloned into p1638 3xMYC. An XhoI – *SFT* 3xMYC-HindIII fragment was subcloned into the pART7 intermediary vector, containing the 35S promoter and the OCS terminator at the XhoI/HindIII site. The final construct was then transferred as a NotI fragment into the ART27 binary vector for transformation. *SP* and *SP5G* were amplified from cDNA with the appropriate primers containing XhoI or SalI at the 5’ and SmaI at the 3’ end (as listed in Suppl. Table 9). *CrFT* was synthesized by GenScript and the *GinFT* cDNA was gift from E. Brenner at NYU. Both were then amplified as the others. The *SFT* in the pART27-35S::*SFT*3XMYC::OCS was then substituted by the appropriate gene using the XhoI and SmaI restriction enzymes. The XhoI-*SFT*-Flag-HindIII fragment was synthesized and cloned into pUC57 by GENEWIZ Inc. A XhoI-SFT-Flag-HindII-SmaI fragment from pUC57 was used to substitute *FT* in the pK2GW7-pHS::*FT*::35S terminator (a gift from Ove Nillson), or cloned into pART7 and pART27 to form pArt27:35S::*SFT*-Flag::OCS. The *SFT* gene was then substituted with the *MIF2* gene that was amplified from a former construct including 43bp upstream to atg. *TFUL2* was cloned from cDNA with the appropriate primers and subcloned into pBS. A SalI-BamHI fragment was cloned into pART7 and then as a Not I fragment into pART27. pPZP212-*pSFT::SFT:*:NOS was cloned by subcloning the 2200bp of p*SFT* and *SFT*::NOS from pPZP111-p*SFT*::GUS::NOS and pCGN1548-35S::*SFT*::NOS, respectively, into pBS. The *pSFT::TFT*::NOS fragment was then cloned to the binary vector pPZP212. The 2800pb of the tomato p*APL* was subcloned into pBS in two fragments: 542bp of the promotor 3’ end was synthesized by GENEWIZ Inc. with StyI at the 5’. The rest was amplified from genomic DNA with appropriate primers (see Suppl. Table 9). The entire promotor was then cloned into the pART27-GUS::OCS cassette that was constructed from pPZP111-p*SFT*::GUS::NOS, pART7 and pART27. Tomato p*CEAS4* was amplified from genomic DNA with appropriate primers (see Suppl. Table 9) and subcloned into pBS. The promotor was then cloned into the pART27-GUS::OCS cassette.

### Construction of CRISPR/Cas9-induced mutant plants

Generation of transgenic plants for CRISPR/Cas9 mutagenesis was performed as per a published protocol (Brooks et al., 2014). Vectors were assembled using the Golden Gate cloning system (Werner et al., 2012). The sgRNAs were cloned downstream of the *At* U6 promoter; sgRNA sequences are listed in Suppl. Table 9. Binary vectors were transformed into M82 and transgenics were genotyped for induced lesions using forward and reverse primers flanking the sgRNA target sites. Next-generation plants carrying mutant alleles and lacking the transgene were used for detailed analyses.

### Sample preparation for NGS analysis

Independent duplicate stem samples consisted of 5 mid-segments of internodes harvested in duplicates 2 h after dawn. Total RNA was extracted using the RNeasy Mini Kit (Qiagen), and treated with DNase I using the RNase-Free DNase set (Qiagen). RNA samples were processed for sequencing, quality control, and differential expression analysis at the Technion Genome Center. Libraries of single 50 bp reads were prepared using the Ilumina TruSeq RNA library preparation kit v2. The barcoded libraries were sequenced on Ilumina Hiseq 2500, six samples per lane (~30,000,000 reads). If an experiment included more than 6 samples, each sample was divided equally, in all the lanes. Library quality control was conducted using FASTQC, version 0.11.5. Sequences were trimmed by quality using trim galore and aligned to the tomato ITAG version 2.5 genome (ftp://ftp.sgn.cornell.edu/genomes/Solanum_lycopersicum/annotation/ITAG2.5_release), using Tophat2, version 2.1.0 (uses Bowtie2 version 2.2.6). Raw gene expression levels were counted by HTseq-count, version 0.6.1 and then normalized by DESeq2 R package, version 1.14.1. Data analysis, gene annotations, and the corresponding A.t. codes, if available, were obtained from the solgenomics ITAG 2.5 release. All Downstream analyses were conducted in R with DESeq2 (Love et al., 2014) as follows: raw data were filtered by independent filtering except when otherwise specified. Count normalization was performed by DESeq2 default. To identify differentially expressed genes (DE/DEG) differing between two conditions (contrast) in the various comparisons described below, we used the Wald test, as implemented in DESeq2. We used padj for adjusted p-value, FC for expression fold change and |FC| for the absolute value of FC.

### NGS analysis

#### Figure1

One factor design (genotype) was used, with three levels (WT, *sft^7187^* and *pSFT:SFT*). Three contrasts were tested: WT vs. *pSFT:SFT*, *sft^7187^* vs. *pSFT:SFT* and *sft^7187^* vs. WT. Genes that significantly differed between conditions were defined as those with padj<0.1 and |FC>2|.

#### Figure 2

Graft – A two-factor design was used, with genotype (WT, *sft*) and induction (homograft, heterograft) as main factors. Genes that significantly differed between homograft and heterograft were defined as those with padj<0.1.

Heat map – The replicate means of the 593 genes showing significant differences between homografts and heterografts (see Suppl. Table 1 for the normalized counts of the 593 genes). Pearson’s correlation as distance matrix and complete agglomeration were used to construct a tree. The tree was then cut into eight clusters and a heatmap was plotted.

Punnett grid (PG) - All 20,286 expressed genes in the recipients WT//WT*, sft//sft and SFT//sft* were sorted into 9 cohorts based solely on their expression trajectories. The expressed genes in each tripartite comparison were defined first by two bipartite genotypic comparisons WT//WT vs *sft//sft* and *sft//sft vs SFT//sft* and subsequently by 3 possible response categories: Up (U), Down (D) and No change (N), demanding a modest |FC≥1.5| as the cutoff.

Cohort DU - genes downregulated in *sft* of the *sft//sft* homograft but upregulated at least 1.5 fold in comparison to *sft* of the *35S SFT//sft* heterograft, is enriched for the florigen–regulated genes analysed in Fig 2C (101/593, p< 5.4*10^−70^ hypergeometric) and for SCWB genes (54/ 365, p< 4.4*10^−26^).

#### Figure 3

Heat induced *FT* –one factor design was used to test the contrast between two factors: heat induced FT and all other conditions (*FT*, heat shock and no added genotype treatment) and genes that significantly differed between conditions were defined as those with padj<0.1 and |FC>1.8|.

#### Figure 4 and 5

*GinFT* and *TFUL2* – two-factor design with genotype and developmental stage as main factors and their interaction. To compute the model, the log-ratio test was used and differential expression was calculated using the Wald test with the specific contrasts. Genes that significantly differed between conditions were defined as those with padj<0.1 and |FC>2|. GO enrichment –DAVID enrichment of biological processes (BP) was applied to all DEGs with known Arabidopsis homologues. Overlap between gene lists – where appropriate, the significance of overlap between lists of genes was tested using the hypergeometric tests.

### qRT-PCR experiments

Total RNA was extracted using the Plant RNA mini kit (Biomiga) and treated with the RNase-Free DNase Set (QIAGEN), according to the manufacturer’s instructions. 500 ng to 1 µg of total RNA was used for cDNA synthesis using the qScript cDNA Synthesis Kit (Quanta). All primer sequences are presented in Suppl. Table 9.

### Histological procedures

Handmade sections of tomato stems were cleared with 5:1 ethanol acetic acid, stained in 0.05% TBO in 20% CaCl_2_ for 2 min, washed in 10% ethanol and kept in 20% CaCl_2_. Images were acquired either by the Nikon eclips e600 microscope equipped with Invenio 3SII camera and Delltapix insight software, or with an Olympus SZX16 binocular microscope equipped with a cooled colour CCD camera (Olympus (DP72) and <u>CellSens</u> software (Olympus).

GUS staining was carried out as previously described (Lifschitz et al., 2006). Briefly, handmade sections of tomato stems were fixed in 90% acetone for 20 min at 4°C, equilibrated in GUS buffer (5mM K_3_Fe(CN)_6_, 5mM K_4_Fe(CN)_6_, 100mM PO_4_ buffer, 1% Triton, 1mM EDTA) for 30 min and stained in GUS buffer contained 25mg/ml X-gluc, at 37°C, for 2-18 h.

### Creating a core list of cell wall genes in tomato

A list of 2765 tomato secondary and primary cell wall genes (SCW and PCW) was compiled andcurated (Suppl. Table 8) using the following sources (Hussay et al, 2013, Caffal and Mahonen 2009). We next screened the list of 180 TFs shown by Taylor-Teeples 2015, to bind regulatory sequences of SCW genes in arabidopsis roots, and added to our list 65 TFs that showed differential expression in at least one of our RNA Seq. experiments. We categorized the genes as SCW, PCW or both based on the above sources. The number of genes in each category and those express in our stem RNA-seq data are shown it Suppl. Table 10.

**Suppl. Table 10.** Categorized CW genes

| Suppl. Table 10 : Categorized CW genes |  |  |
| --- | --- | --- |
|  | Total | Expressed in stem |
| All | 34728 | 28728 |
| Cell Wall | 2675 | 2440 |
| PCW | 273 | 238 |
| SCW & PCW | 24 | 24 |
| SCW | 498 | 468 |

### Western blot analysis

Protein sample concentrations were assessed via the Bradford assay. Equal amounts of protein were prepared for loading with sample buffer (0.1mM Tris-HCl pH 6.8, 2% SDS, 4% glycerol, 400mM dithiothreitol, 0.01% bromophenol blue), heated for 5 min at 100 °C and centrifuged for 1 min. The supernatant separated on a 12.5% SDS-PAGE gel (1 h, in 25mM Tris-HCl, 192 mM glycine buffer, 0.1% SDS). Proteins were transferred to a nitrocellulose membrane (Protran) via wet transfer at 4 °C in a 25mM Tris-HCl, 192 mM glycine 20% methanol and 0.03% SDS buffer. Membranes were blocked with 1% skim milk, for 1 h, at RT, and incubated with a primary antibody overnight, at 4 °C. Membranes were then washed 3 times with TBST (20mM Tris-HCl pH7.6, 150mM NaCl, 0.1% Tween-20) and incubated for 1 h, at RT, with the secondary antibody conjugated to horseradish peroxidase. Then, membranes were washed 3 times with TBST and an ECL reaction was performed. The protein signal was visualized with a Biorad-Chemidoc analyzer.

### Enrichment for the mobile MYC-tagged CETS proteins

Leaves or stems from recipient grafts were ground in liquid nitrogen and stored at −80 °C. For SFT-3XMYC, the tissue powder was extracted in 5 volumes of Mg/NP40 extraction buffer (Kim et al., 2001), mixed with 4 volumes of 90% cold acetone, and stored at −20°C for 1h. The acetone mix was briefly precipitated, the pellet was resuspended in HEPES-Sorbitol buffer, and then precipitated again at 100,000 g for 1h. The final supernatant was subjected to ammonium sulfate (AS) fractionation as detailed in the Supplemental Methods. Enrichment for other CETS proteins was performed without the prior acetone purification. Tissue powder was homogenized in Mg/NP40 (Kim et al., 2001) at a 1:5 w/v tissue:buffer ratio. [0.5M Tris-HCl, pH 8.3, 2% v/v NP-40, 20mM MgCl2, 2% v/v β-mercaptoethanol, 1mM phenylmethylsulfonylfluoride (PMSF) and 1% w/v polyvinylpolypyrrolidone (PVPP)], for 15 min 12,000 g and subjected to AS fractionation (supplemental methods).

### Purification of CETS proteins for MS analysis

Plant tissue (10 g) was extracted with Mg/NP40 followed by AS fractionation, as above, with the exception of SP-3xMYC which was subjected to a 20-50% fractionation. The pellets were suspended in 2 ml 50mM Tris-HCl, pH8, 150mM NaCl, filtered through a 0.45 µm filter (Minisart Sartorius^®^), and then purified on anti-MYC columns. Anti-MYC column preparation. General Reference Protein A beads, for a 50 µl bed volume, were briefly centrifuged at 3000 rpm and pellets were suspended in 0.5ml NaBr buffer [50mM NaBr pH 9, 3M NaCl]. anti-MYC (Hybridoma purified, see suppl M&M; 50 µg) in 0.5 ml NaBr buffer were combined with the beads in a test tube and rotated for 1 h, at RT. After centrifugation at 10,000g for 30 sec, beads were washed with NaBr buffer, centrifuged, and resuspended in 2 ml 200 mM NaBr pH 9, 3M NaCl. Dimethyl pimelimidate (DMP) was added to bring the same to a 20 mM concentration for crosslinking and the test tube was rotated for 30 min at RT. After centrifugation and resuspension in 0.2M ethanolamine, pH 8, the test tube was rotated for 2 h, at RT. The test tube was then centrifuged and beads resuspended in PBS, pH 7.4, the crosslinked beads were placed in a tip column (TopTip Empty Glygen^®^) and stored at with 0.02% Na-azide, at 4 °C until use.

### Immunoaffinity purification of MYC-tagged proteins for MS

An anti-MYC column was washed with 1 column volume of acetic acid, pH 3, 10 column volumes of Tris-HCl 100mM, pH 8 and 2 column volumes of 50 mM Tris-HCl, pH8, 150 mM NaCl. Samples (2ml) were loaded, washed with 3 volumes of 50 mM Tris-HCl, pH 8, 150 mM NaCl and eluted in 10 fractions of 50 µl acetic acid 0.1 N, pH 3, 8 µl 1M Tris-HCl, pH9 were added per fraction for neutralization. Samples of the fractions were Western blotted and strongly reacting fractions were used for mass spectrometry analysis.

### LC-MS/MS

Samples were either trypsinized or chymotrypsinized and peptides were analyzed by LC-MS/MS, performed with an OrbitrapXL mass spectrometer (Thermo). The data were analyzed using the Sequest 3.31 software vs the Solanum lycopersicon section of the NCBI-NR database and vs. the specific protein sequences. SP-3xMYC results were further validated via higher-energy collisional dissociation (HCD) fragmentation (see suppl Methods for further details). All analyses were carried out at the Smoler Proteomics Center, Department of Biology, Technion - Israel Institute of Technology Haifa Israel.

## Supporting information

## Acknowledgments

Funding-This work was supported by Israel Science Foundation Research (ISF) and Russell Berrie Nanotechnology Institute grants to E.L, by ISF and BARD grants to Y.E and by a joint BIKURA grant to E.L. and Y.E.

## Author contributions

T. E-R, A S-K, and N. S. performed experiments, T. E-R organized the cloning and the RNA-seq experiments, G.H performed the RNA-seq statistical analysis and organized the graphical illustrations together with T. E-R, A. S-K and E.L. R.A. contributed his immense anatomical experience and insights. Y.E. provided an excellent intellectual input, contributed tomato lines and the CRISPR –edited lines. E.L conceived and led the project, conducted the grafting, histological, and genetic experiments and wrote the paper with input from the other authors.

