## Supplementary material for "The Flowering Hormone Florigen Accelerates Secondary Cell Wall Biogenesis to Harmonize Vascular Maturation with Reproductive Development"

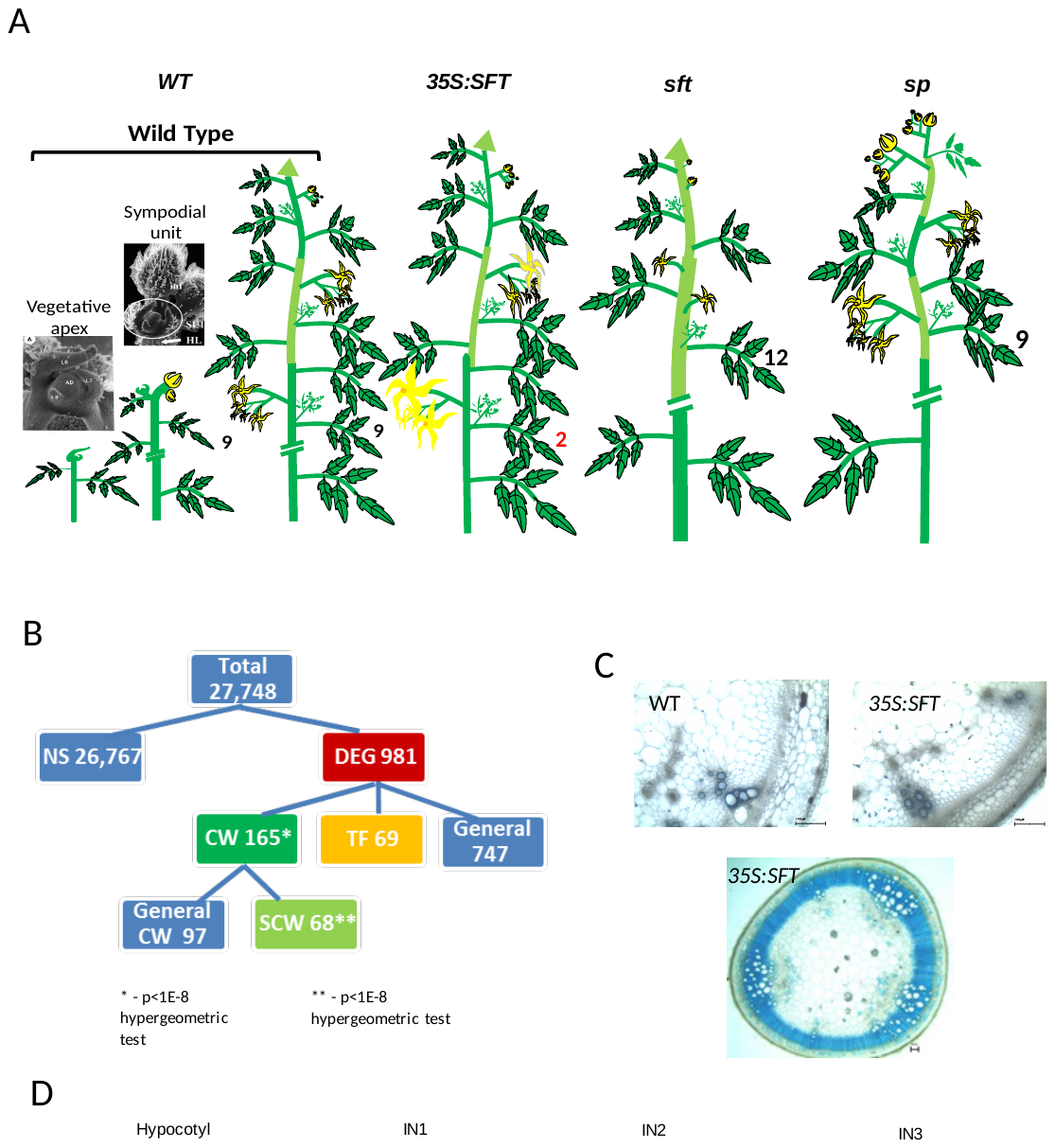

Suppl. Fig. 1 High expression of *SFT* (florigen) accelerate SCWB in the tomato stem

(**A**)The sympodial growth habit of tomato in sympodial plants, the primary apical bud is terminated by differentiating organs, typically inflorescences. Subsequently the upper most axillary bud, called sympodial, is released and eventually displaces the terminal inflorescence sideways. The reiteration of this process results in a compound sympodial shoot. In tomato, depending on the integral light dose, the primary shoot is terminated after 7-12 leaves. The first and subsequent sympodial units are terminated by an inflorescence after forming 3 leaves. Therefore, flowering is synonymous with termination. Termination of vegetative apices, being primary or sympodial, is required for the activation of the next sympodial cycle. In *sft* mutant plants, primary flowering is delayed by ca. five leaves, the terminating inflorescence consists of a single flower inflorescence with a leafy ad-axial sepal and additional leaves instead of flowers. The vegetative inflorescence functions as a regular shoot apical meristem thus arresting differentiation and disrupting the regular sympodial cycle. Over expression of *SFT* induces flowering after only three leaves and in addition delays the first sympodial bud which is later released as a regular lateral branch. This first ‘lateral’ branch forms a regular sympodial shoot with the typical reiterated 3 node sympodial units (Shalit et al., 2009). Overexpression of *SELF PRUNING* (*SP*) delays flowering, thus serving as a florigen antagonist but unlike *SFT* the termination of the primary shoot is not affected. The sympodial pattern is also normal but the sympodial units in *self pruning* plants become progressively simple until the shoot is terminated with two consecutive inflorescences (Pnueli et al., 1998). (**B**) Summary of the classification of the transcriptome profiling of tomato stems. Functional classification of 27,748 expressed genes- * padj<0.1 and absolute fold change ≥2.RNA profiles of the 3^rd^ internodes of *pSFT:SFT*, WT and *sft* plants. Genes that significantly differ (padj<0.1 and absolute fold change >=2) between WT vs *pSFT:SFT* or *sft^7187^*vs *pSFT:SFT*. (**C**) Top: Cross sections of stems from 20 day old WT and *35S: SFT* plants. The *35S: SFT* plants developed stage 10 inflorescences while only floral primordia differentiated in WT plants. No secondary growth was observed at that time in the IF regions of both phenotypes. -Bottom, a cross section of a mature *35S:SFT* plant stem. TBO staining for lignin. (**D**) Expression of the pCesA-GUS reporter gene in hypocotyl and sequential stem internodes of a 3 week old WT plant. GUS staining.

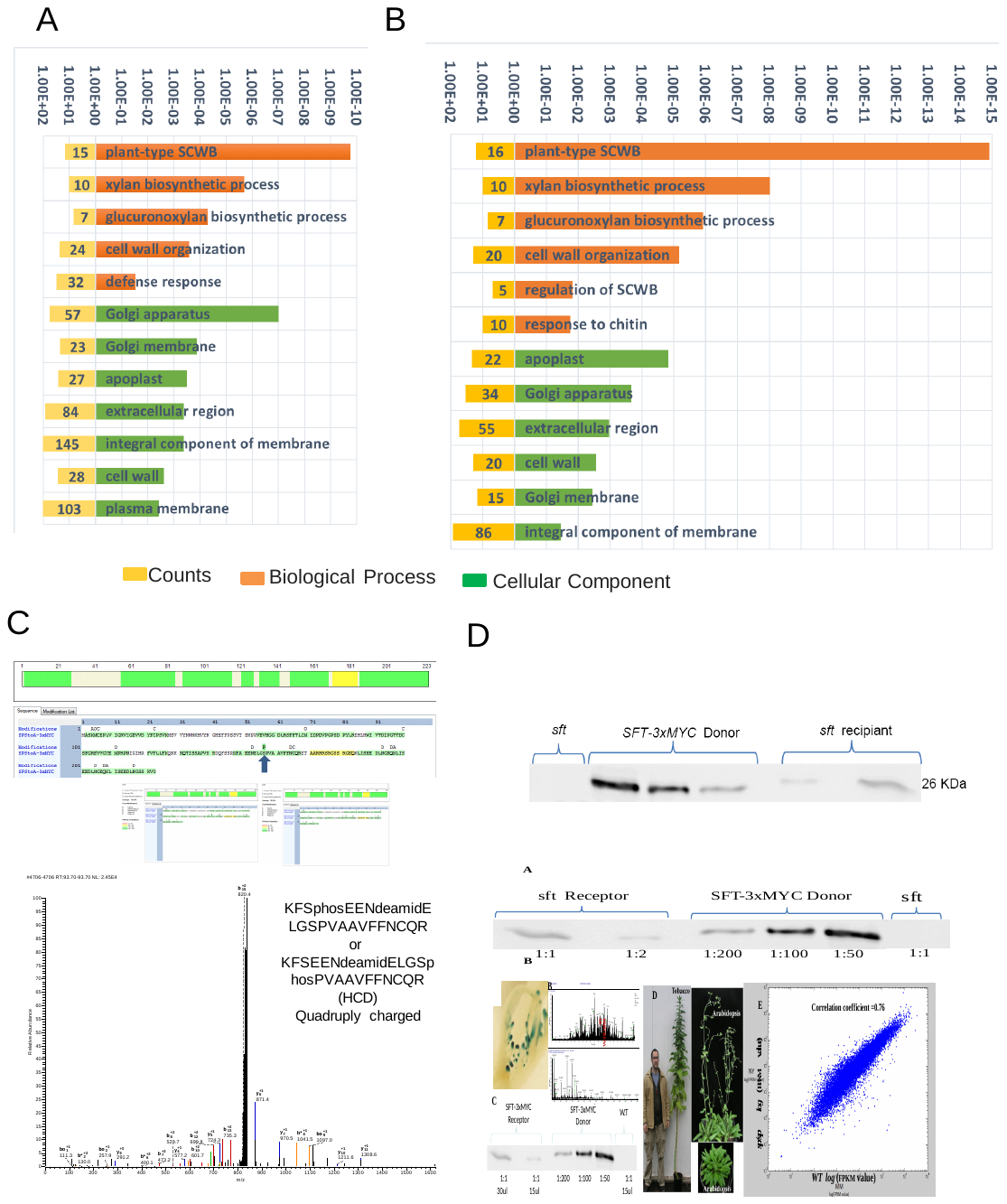

Suppl. Fig. 2 The m-florigen

(**A**) Functional classifications (Biological Processes and Cellular Components) of the 593 m-florigen-responsive genes in the grafting experiment. The 12 top categories are shown. (**B**) Functional characterization of the 365 genes of cohort DU of PG1 (Fig2E). (**C**) Affinity enriched SP-MYC is phosphorylated serine residue at position 157. (**D**) The mobile SFT-MYC protein is free of major post-translation modifications. Top- calibration of the affinity-purified mobile SFT-MYC protein.

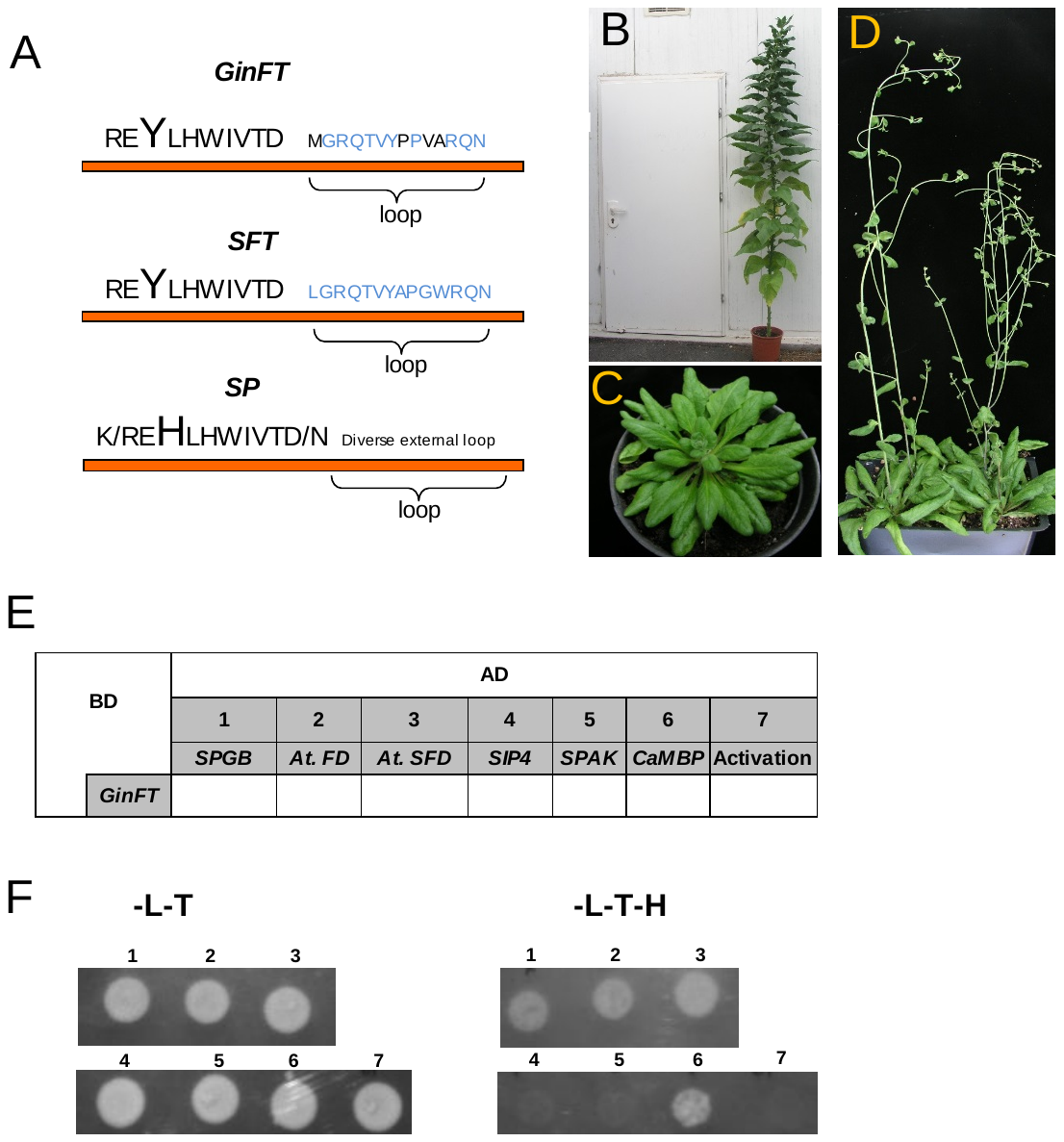

Suppl. Fig. 4 GinFT interactions

(**A**) Amino acids sequence alignment of the GinFT, SFT and SP polypeptides. In tomato *GinFT* functions as a florigen antagonist but carries Y88, which is conserved in the FT (SFT) clade. In contrast GinFT features a divergent external loop as typical to florigen antagonists of the TFL (SP) clade of CETS proteins. (**B-D**) Overexpression of *GinFT* delays flowering in Arabidopsis and tobacco. (**B**) A transgenic *35S:GinFT* tobacco plant. Such plants form more than 80 leaves (vs. 24 of WT) before terminating with a smaller but normal inflorescence. (**C-D**) *35S:GinFT* Arabidopsis plants grown under long days. The term “florigen antagonist” refers to homologs of florigen, genes that suppress flowering when overexpressed but enhance it when inactivated.

(**E** -**F**) GinFT interacts with tomato and Arabidopsis FD transcription factors. These results were extracted from an experiment performed beforehand for other purposes (**F**) and were also summarized in a table (**E**). The original plates are available upon request. The numbers of the bottom colonies refer to those in the table above.

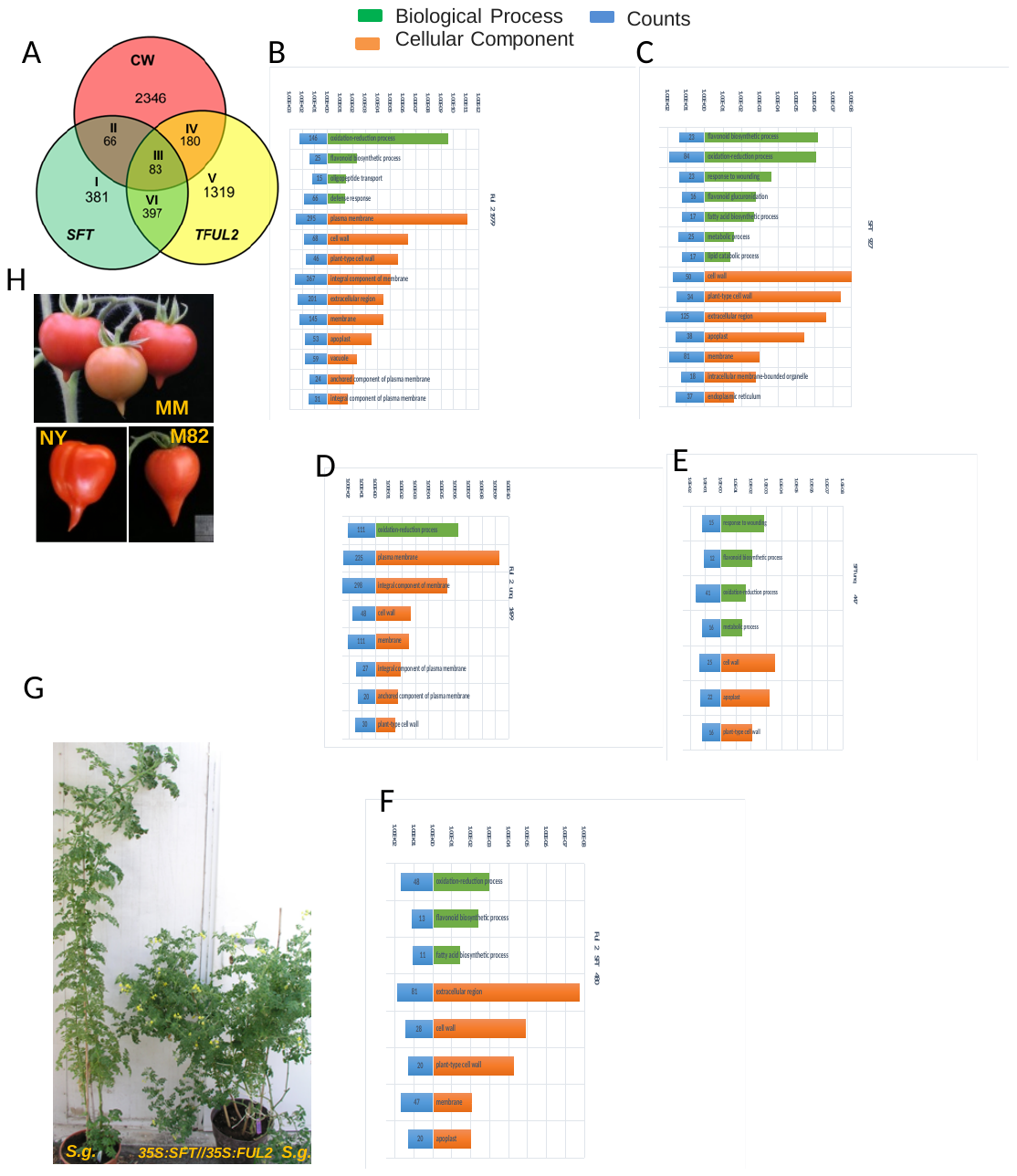

**Suppl. Fig. 5 Common and independent biological functions of *SFT* and *TFUL2***

(**A-F**), A detailed functional analysis of *SFT* and *TFUL2* regulated genes as classified in Venn diagram in Fig. 5J. (**A**)- A Venn diagram, duplicating text Fig. 5 J. (**B**) GO terms of *TFUL2*-specific regulated genes (Class III+IV+V+VI in (A)). (**C**) All *SFT* regulated genes (class I+II+ III+VI in (A)). (**D**)*TFUL2*- specific regulated genes (class V&VI in (A)). (**E**) *SFT*-specific regulated genes (class I+II in (A)). (**F**) genes regulated by both *SFT* and *TFUL2* (class II+VI in(A)). (**G**) Tomato donor of florigen induces flowering in *S.galapagense* recipient plants overexpressing *TFUL2*. Both plants were grown under long days. (**H**) Abundant expression of *TFUL2* induces pointed beaked fruits in all backgrounds. Similarly, overexpression of four other *TFUL* like genes; *AtAP1*, *MACROCALYX*, *TFUL1* and *AP67* (see Table S1) induced slender stems and beaked fruits in all 3 backgrounds.

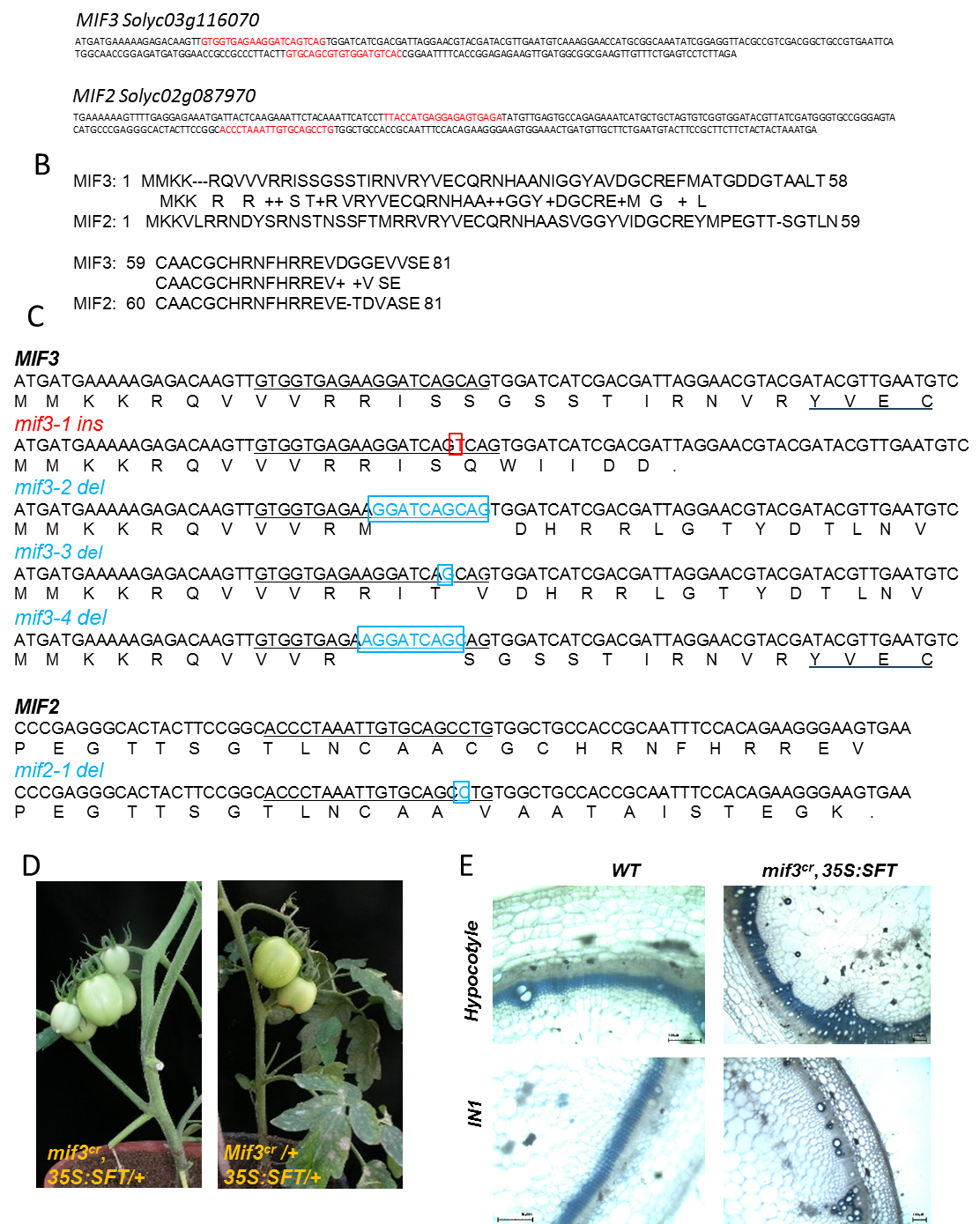

Suppl. Fig. 6 Genome editing of the *MIF2* and *MIF 3* genes

(**A**) RNA guides (Red) for the tomato *MIF2* and *MIF3* genes. (**B**) Amino acid alignment for MIF2 and MIF3 polypeptides. (**C**) CRISPR – edited alleles of *MIF3* Top and *MIF2*, Bottom. Insertions, Red and Deletions, Blue. (**D**) The effect of high SFT on stem sizes of mature *mif3 ^cr1^* plants. Fruit bearing stems of *mif3^cr1^ 35S:SFT/+* and *mif3 ^cr1^/+ 35S:SFT/+* plants. Note the normal radial size of *mif3 ^cr1^ 35S:SFT/+* stems. (**E**) Comparison of IF vascular differentiation in hypocotyls and stems of 32 day old WT and *mif3^cr1^ 35S:SFT* plants. TBO staining for lignin.

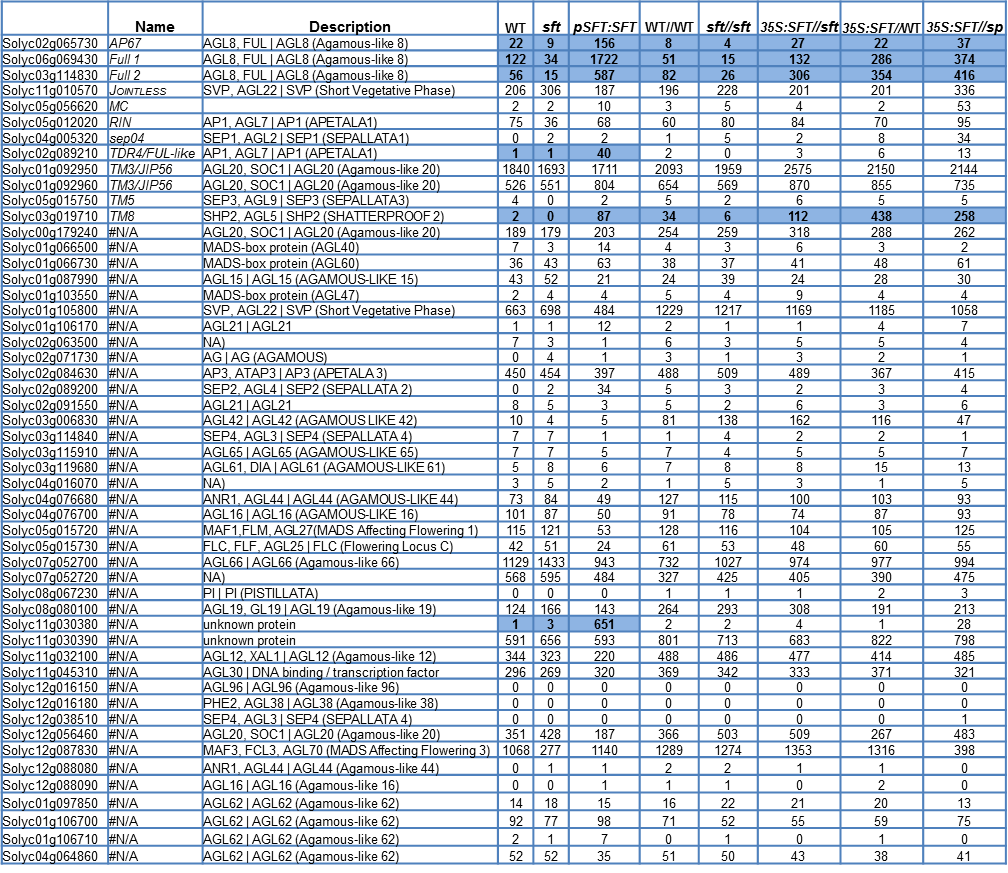

Suppl. Table 2 MADS genes

The expression profiles of MADS genes in the graft experiments.

Suppl. Data (available upon request)

Table S1: WT vs p *SFT-SFT*

Table S3: Graft

Table S4: DU

Table S5: Heat Shock

Table S6: *GINFT*

Table S7: *TFUL2-SFT* Venn

Table S8: Cell wall list

Table 9: primers for RTPCR and Cloning

**Table S1: WT vs p *SFT-SFT***

List of differentially expressed genes between *WT* and *pSFT-SFT*

is provided as a separate file in MS Excel (.xls) Format. Only genes showing larger

than two-fold change and *P* value < 0.1 are listed. Additional columns show the sum of mean normalized Deseq2 expression values for WT, *sft* mutant and *pSFT-SFT* overexpression (column B-D), whether a particular gene belongs to a Transcription Factor (TF) family (column E), Cell wall (column F), primary or secondary cell wall (column G), description (column H & J) the closest *Arabidopsis thaliana* homolog (column J). The remaining sheets show only TF and only SCW.

**Table S3: Graft**

List of differentially expressed genes between grafts is provided as a separate file in MS Excel (.xls) Format. Only genes showing P value < 0.1 are listed (see M&M for further details). Additional columns show the sum of mean normalized Deseq2 expression values for *WT//WT, sft//sft, 35S:SFT//sft, 35S:SFT//WT, 35S:SFT//SP*, (column B-F), whether a particular gene belongs to a Transcription Factor (TF) family (column G), Cell wall (column H), primary or secondary cell wall (column I), description (column J& K) the closest *Arabidopsis thaliana* homolog (column L) and the cluster in the heatmap (column M).

**Table S4: DU**

List of differentially expressed genes between grafts is provided as a separate file in MS Excel (.xls) Format. Only genes downregulated in *sft//sft* homograft but upregulated at least 1.5 fold in comparison to *35S SFT//sft* heterograft are listed (see M&M for further details). Additional columns show the sum of mean normalized Deseq2 expression values for *WT//WT, sft//sft, 35S:SFT//sft, 35S:SFT//WT, 35S:SFT//SP*, (column B-F), whether a particular gene belongs to a Transcription Factor (TF) family (column G), Cell wall (column H), primary or secondary cell wall (column I), description (column J& K) the closest *Arabidopsis thaliana* homolog (column L).

**Table S5: Heat Shock**

List of differentially expressed genes induced by heat shock is provided as a separate file in MS Excel (.xls) Format. Only genes showing *P* value < 0.1 and |FC|>1.8 are listed. Additional columns show the sum of mean normalized Deseq2 expression values for treated *sft pHS:FT*

untreated *sft pHS:FT,* treated *sft* and untreated *sft* (column B-E), whether a particular gene belongs to a Transcription Factor (TF) family (column F), Cell wall (column G) , primary or secondary cell wall (column H), description (column I&J) and the closest *Arabidopsis thaliana* homolog (column K).

**Table S6: Gin*FT***

List of differentially expressed genes between *WT* and *35S:GinFT*  is provided as a separate file in MS Excel (.xls) Format. Only genes showing |FC|>2 and *P* value < 0.1 are listed. Additional columns show the sum of mean normalized Deseq2 expression values for WT and *GinFT* (column B-C), whether a particular gene belongs to a Transcription Factor (TF) family (column D), Cell wall (column E) , primary or secondary cell wall (column F), description (column G & H) and the closest *Arabidopsis thaliana* homolog (column I).

**Table S7: *TFUL2- SFT* Venn**

List of differentially expressed genes between *WT* and *35S:TFUL2 or WT* and *35S:SFT* is provided as a separate file in MS Excel (.xls) Format. Only genes showing |FC|>2 and *P* value < 0.1 are listed. Additional columns show the sum of mean normalized Deseq2 expression values for WT, *35S:TFUL2* and *35S:SFT* (column B-D), whether a particular gene belongs to a Transcription Factor (TF) family (column E), Cell wall (column F) primary or secondary cell wall (column G), description (column H&I) and the closest *Arabidopsis thaliana* homolog (column J). In the first sheet all genes showing DEG between WT and *35S:SFT,* In the second sheet all genes showing DEG between WT and *35S:TFUL2.* In the third sheet only genes showing DEG between WT and *35S:SFT* but not between WT and *35S:TFUL2* )*SFT* unique). In the fourth sheet only genes showing DEG between WT and *35S:TFUL2* but not between WT and *35S:SFT* *(TFUL2* unique). In the fifth sheet, genes regulated by both *SFT* and *TFUL2.*

**Table S8: Cell wall list**

List of 2675 genes that were defined as a cell wall genes as described in M&M. Additional columns show whether a particular gene belongs to a Transcription Factor (TF) family (column B), Cell wall (column C), primary or secondary cell wall (column D), description (column E&F), and the closest *Arabidopsis thaliana* homolog (column G), the reference and the specific description for a given gene (column H & I). The last column (J) indicate if the gene expressed (at least over 1 read) in our data.

**Table S9: primers for qRT-PCR and Cloning**

**Supplemental Methods**

Anti-MYC purification

An anti–MYC hybridoma was grown in standard cRPMI (10% FCS + Glu + pen/strep). Cells were transferred to a starvation medium (50 ml cRPMI with biogrowth instead of FCS) and incubated for 5-7 days. Medium was filtered and diluted 1:1 with binding buffer (0.02M NaH2PO4, 0.15M NaCl, pH8). A Protein A column with a 0.5 ml bed volume was prepared according to the manufacturer's instructions. A Sepharose 4B Fast flow beads, Sigma®). Diluted cell culture was loaded onto the column via a peristaltic pump at a flow rate of 15-50 ml/h, at 4 °C. Beads were washed with binding buffer and eluted in 5 fractions of 250 μl of citrate buffer (0.2M Na2HPO4, 0.1M citric acid, pH 4.5-5 ) and immediately neutralized with 80 μl Tris-HCl, pH9. Fractions with a concentration of over 0.3 mg/ml were pooled and dialyzed overnight against PBS, pH7.4.

Ammonium sulfate precipitation

Cold (4ºC) saturated AS was added to protein extracts, to the percent of saturation determined as optimal. The solution was mixed well, chilled 30 min on ice and centrifuged 15min, 12,000g. Supernatant was then subjected, if required, to additional AS fractionations. Pellets were re-suspended in HEPES-sorbitol buffer [50mM HEPES, 10% sorbitol, 150mM NaCl, 1mM EDTA, 5mM MgCl2]. AS fractions used for Western blot were: SFT-3xMYC 35-55%, SP-3xMYC 0-20%, SP5G-3xMYC 30-60%, SP2G-3xMYC 30-60%, GnKFT-3xMYC 30-50%. Insoluble material in the SP-3xMYC pellet was extracted by boiling in x1 SDS sample buffer, after centrifugation at 12,000g for 15 min, the supernatant was added to the soluble material.

LC-MS/MS analysis

Identification results were filtered according to the cross-correlation score (Xcore value), mass accuracy and the probability and presented as a combined report. Probability (P) displays the probability of finding a match as good as or better than the observed match by chance. The value displayed for the protein is the probability of the best peptide match (the peptide with the lowest score). A peptide was considered as high quality if receiving a SequestXcore of 1.2 for singly charged peptides, 2.0 for doubly charged peptides and 2.5 for triply charged peptides
